# Post-translational polymodification of *β*1 tubulin regulates motor protein localisation in platelet production and function

**DOI:** 10.1101/595868

**Authors:** Abdullah O. Khan, Alexandre Slater, Annabel Maclachlan, Phillip L.R. Nicolson, Jeremy A. Pike, Jasmeet S. Reyat, Jack Yule, Rachel Stapley, Steven G. Thomas, Neil V. Morgan

## Abstract

In specialised cells, the expression of specific tubulin isoforms and their subsequent post-translational modifications drive and coordinate unique morphologies and behaviours. The mechanisms by which *β*1 tubulin, the platelet and megakaryocyte (MK) lineage restricted tubulin isoform, drives platelet production and function remains poorly understood. We investigated the roles of two key post-translational polymodifications (polyglutamylation and polyglycylation) on these processes using a cohort of thrombocytopenic patients, human induced pluripotent stem cell (iPSC) derived MKs, and healthy human donor platelets. We find distinct patterns of polymodification in MKs and platelets, mediated by the antagonistic activities of the cell specific expression of Tubulin Tyrosine Ligase Like (TTLLs) and Cytosolic Carboxypeptidae (CCP) enzymes. The resulting microtubule patterning spatially regulates motor proteins to drive proplatelet formation in megakaryocytes, and the cytoskeletal reorganisation required for thrombus formation. This work is the first to show a reversible system of polymodification by which different cell specific functions are achieved.

**Key Points:**

- The platelet specific *β*1 tubulin (encoded by *TUBB1*) is polymodified (polyglutamylated and polyglycylated) in platelet producing iPSC-derived megakaryocytes (MKs), this patterning spatially regulates motor proteins, and its disruption in *TUBB1* patient variants results in a loss of platelet production.
- A system of reversible polymodifications mediated through the graded expression of modifying enzymes (TTLLs and CCPs) throughout MK maturation is required for proplatelet formation and subsequent platelet function.

## Introduction

Microtubules are large, cytoskeletal filaments vital to a host of critical functions including cell division, signalling, cargo transport, motility, and function(1–3). Despite their ubiquitous expression and high structural conservation, microtubules also drive unique morphologies and functions in specialist cell types like ciliated cells, spermatozoa, and neurons(4, 5). The question of how filaments expressed in every cell in the body can facilitate complex and highly unique behaviours like neurotransmitter release and retinal organisation has been addressed by the *tubulin code*. This is a paradigm which accounts for the specialisation of microtubules and their organisation by describing a mechanism in which particular cells express lineage restricted isoforms of tubulin. These cell specific isoforms are then subject to a series of post-translational modifications (PTMs) which alter the mechanical properties of microtubules, and their capacity to recruit accessory proteins (e.g. motor proteins)(1–3, 5).

A host of PTMs have been reported in a range of cell types, including (but not limited to) tyrosination, acetylation, glutamylation, glycylation, and phosphorylation. The loss of specific tubulin PTMs, either through the aberrant expression of tubulin isoforms or the loss of effecting or reversing enzymes, has been linked to disease and dysfunction in motile and non-motile cilia (including respiratory cilia, retinal cells), spermatogenesis, muscular disorders, and neurological development (1, 5–11). Despite an increasing understanding of the importance of tubulin PTMs in disease, the role of the tubulin code in the generation of blood platelets from their progenitors, megakaryocytes (MKs) remains poorly understood.

Platelets are the smallest component of peripheral blood, and circulate as anucleate cells with an archetypal discoid shape maintained by a microtubule marginal band(12, 13). Antagonistic motor proteins maintain the resting state of the marginal band, and during platelet activation a motor dependent mechanism results in sliding which extends the marginal band and causes the transition to a spherical shape (13–15).

Conversely megakaryocytes are the largest and rarest haematopoeitic cell of the bone marrow. These cells are characteristically large, polyploid cells with unique morphological structures (e.g. the invaginated membrane system (IMS)) required to facilitate the production of thousands of blood platelets and package within them the required pro-thrombotic factors(15, 16). MKs form long, beaded extensions into the lumen of bone marrow sinusoids - where these proplatelet extensions then experience fission under the flow of sinusoidal blood vessels which results in the release of barbell shaped pre-platelets and platelets into the blood stream(16).

Both MKs and platelets express a lineage restricted isoform of *β* tubulin (*β*1 tubulin) encoded by the gene *TUBB1*(17). In humans, *TUBB1* mutations have been shown to result in impaired platelet production, with a resulting macrothrombocytopenia(18, 19). More recently, a C-terminal truncation of *β*1 tubulin has been shown to cause a macrothrombocytopenia, suggesting that C-terminal modifications may be drivers of protein function and causative of the disease phenotype observed (20).

While the loss of *TUBB1* is known to result in macrothrombocytopenia, the mechanisms by which this isoform of tubulin effects the dramatically different cytoskeletal behaviours of platelets and MKs remains poorly understood. In the context of the tubulin code, MKs and platelets present a particularly interesting model. Both cells express *β*1 tubulin, but undergo markedly different cytoskeletal changes. To date, acetylation and tyrosination have been the PTMs primarily reported in MKs and platelets, however neither modification is specific to the C-terminal tail encoded by *TUBB1*. Fiore *et al*. show that a C-terminal truncation of *TUBB1* phenocopies the complete loss of the protein(13, 21). We therefore hypothesise that PTMs specific to the C-terminus of *TUBB1* are required for the complex morphological rearrangements required for both MK and platelet function. The C-terminal tail of *β*1 tubulin is particularly rich in glutamate residues which are often targeted for two key PTMs implicated in human disease. Polyglutamylation and glycylation are PTMs which target glutamate residues on both tubulin subunits (*α* and *β*) and result in the addition of glutamate or glycine residues respectively(1, 2, 22). Interestingly polyglutamylation has been observed in microtubules in centrioles, axenomes, neuronal outgrowths, and mitotic spindles(1, 2).

Thus far polyglycylation has primarily been observed in axenomes, suggesting a role for polyglycylation and polyglutamylation in regulating ciliary function - with important consequences for ciliopathies(1). As these polymodifications target the same substrate, namely glutamate residues in tubulin tails, it has been suggested that these PTMs are competitive. For example, glutamylation is evident on *β* tubulin in post-natal development, but is found on *α*-tubulin in younger neurons(23). There is some debate as to whether these polymodifications negatively regulate one another(24, 25). Mutations in the glutamylases and deglutamylases regulating polymodification can cause male infertility through aberrant spermatogenesis and poor sperm motility, as well as dysfunctions in airway cilia and axonal transport (9, 11, 26–29)

To date polyglycylation has not been reported in MKs or platelets. Recently Van Dijke *et al*. reported on the polyglutamylation of *β*1 tubulin in a CHO cell line engineered to express *TUBB1* downstream of the integrin 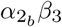, and platelets spread on fibrinogen(30). However, to effectively study and interpret the effects of post-translationally modified tubulin residues a model is required which recapitulates the complex network of regulatory enzymes effecting and reversing PTMs. This is particularly important in light of potential evolutionary divergences in the function of TTLL enzymes implicated by Rogowski *et al*.(25). To interrogate the polymodification of the C-terminal tail of *TUBB1* as a driver of both platelet formation and function, we report on two unrelated families with genetic variants in the C-terminal region of *TUBB1* gene, resulting in a macrothrombocytopenia and bleeding. Existing cell lines poorly emulate the expression profiles and platelet producing capacity of MKs, therefore to develop a representative model we adapt a directed differentiation protocol to generate a large population of proplatelet forming MKs from induced pluripotent stem cells (iPSCs). We apply these approaches to report for the first time the extensive polyglycylation and polyglutamylation of human proplatelet forming MKs. We demonstrate a markedly different pattern of polymodification in resting platelets, which lose tubulin polyglylycylation but maintain polyglutamylation of the marginal band. We report that upon activation, platelets undergo a specific polyglutamylation of the marginal band.

We hypothesise that the distinctive patterning of polymodification in platelets and MKs regulate the localisation of key motor proteins known to drive platelet production and shape change upon platelet activation. To that end, we perform a quantitative mRNA-expression analysis of the 13 known mammalian tubulin tyrosin like ligases (TTLLs) and 6 Cytosolic Carboxypeptidases (CCPs) and report an increase in the expression of specific TTLLs and CCPs in maturing and proplatelet forming MKs. Our *in vitro* work identifies a mechanism by which maturing MKs express a palette of TTLLs and CCPs to reversibly polyglutamylate and polyglycylate *β*1 tubulin to regulate motor protein motility and drive platelet production. Knocking-out *TUBB1* in iPSC-MKs results in a loss of platelet production, and disordered polymodification. In platelets a single polyglutamylase, TTLL7 is expressed to drive the polyglutamylation of the marginal band needed to destabilise this structure for platelet activation and shape change. This study reports on a highly unique mechanism of reversible polymodification which drives the specialist behaviours of both platelets and their progenitors, MKs. Finally we report a novel cause of bleeding due to patient variants in 3 unrelated families which result in a loss-of-function of the tubulin modifying polyglycylase *TTLL10*, suggesting a role for this gene, previously thought to be non-functional in humans, in platelet production.

## Materials and Methods

### Study Approval

Whole blood was obtained for each experiment from healthy volunteers under the University of Birmingham’s ERN 11-0175 license ‘The regulation of activation of platelets’. The Genotyping and Phenotyping of Platelets (GAPP) study was approved by the National Research Ethics Service Committee West Midlands – Edgbaston (REC reference 06/MRE07/36). Participants gave written informed consent in compliance with the Declaration of Helsinki. The GAPP study is included in the National Institute for Health Research Non-Malignant Haematology study portfolio (ID 9858), and registered at ISRCTN (http://www.isrctn.org) as ISRCTN77951167.

### Whole exome sequencing (WES)

To identify the causative mutation in these families we sequenced the whole exome of the affected individuals with the SureSelect human AllExon 50Mb kit (Agilent Technologies) and sequenced on the HiSeq 2500 (Illumina) with 100 bp paired-end reads. The sequences were aligned to the reference genome (hg19 build), with Novoalign (Novocraft Technologies Sdn Bhd). Duplicate reads, resulting from PCR clonality or optical duplicates, and reads mapping to multiple locations were excluded from downstream analysis. Depth and breadth of sequence coverage was calculated with custom scripts and the BedTools package. Single nucleotide substitutions and small insertion deletions were identified and quality filtered within the SamTools software package and in-house software tools. All calls with a read coverage <4 and a phred-scaled SNP quality of <20 were filtered out. Genetic variants were annotated with respect to genes and transcripts with the Annovar tool. Candidate variants were prioritised according to their frequency in the latest population databases where a rare variant was defined as <0.01.

### Platelet preparation

25mL volumes of blood were drawn from volunteers into sodium citrate. PRP (platelet-rich plasma) was generated by centrifugation of samples for 20 minutes at 200g. PRP was further spun to isolate platelets by centrifugation at 1,000g for 10 minutes with prostacyclin (0.1 *µ*g/mL) and ACD. The resulting pellet was suspended in Tyrode’s buffer prepared fresh, and pre-warmed at 37°C (5mM glucose, 1mM MgCl2, 20mM HEPES, 12mM NaHCO3, 2.9 mM KCL, 0.34mM Na2HpO4, 129mM NaCl to a pH of 7.3). This suspension was spun again at 1,000g with prostacyclin at the same concentration before re-suspended to a final concentration of 2 × 10^8^/mL. Platelets were left to rest for 30 minutes at room temperature before any further processing or treatment.

### Platelet spreading

Resting platelets were fixed by preparing platelets at a concentration of 4 × 10^7^/mL and mixing with equal volumes of 10% neutral buffered Formalin in a 15mL falcon tube. This mixture was inverted gently to mix the sample, and left to incubate for 5 minutes before subsequently adding 300*µ*L of the resulting fixed, resting platelets to coverslips coated in Poly-L-Lysine (Sigma). Cells were then spun down at 200g for 10 minutes.

Spreading was performed on human fibrinogen (Plasminogen, von Willebrand Factor and Fibronectin depleted - Enzyme Research Laboratories) and Horm collagen (Takeda). Coverslips were coated overnight at a concentration of 100 *µ*g and 10*µ*g/mL (fibrinogen and collagen) respectively, before blocking for 1 hour in denatured fatty acid free 1% BSA (Life Technologies). Finally coverslips were washed once with PBS before the addition of platelets. Unless otherwise stated (as in the time course experiment), platelets were spread for 45 minutes at 37°C. Fixation for spread platelets was performed in formalin as for resting platelets for 10 minutes.

### Construct cloning and transfection

*β*1-tubulin wild type construct was generated by the gibson assembly (HiFi Kit, New England Biolabs) of a *TubB1* sequence fragment synthesised as a gBlock by Integrated DNA Technologies (IDT) and a C-terminal mApple empty backbone (mApple-C1 was a gift from Michael Davidson (Addgene plasmid # 54631; http://n2t.net/addgene:54631; RRID:Addgene_54631)(31). All mutagenesis experiments were performed using the Q5 site directed mutagenesis kit from New England Biolabs following their supplied protocol. The table in supplementary figureS2 provides the primers and annealing temperatures used to produce the various mutants using the mAPPLE TubB1 plasmid as a template. All primers were designed using the online NEBase changer tool. Transfection was performed using a standard Lipofectamine 3000 protocol in Hek293T cells maintained in complete DMEM as described in the author’s previous work (32, 33).

### Immunofluorescence

After fixation platelets were washed twice with PBS before incubation in 0.1% Triton X-100 for 5 minutes. The subsequently permeabilised cells were washed twice with PBS before blocking in 2% Goat serum (Life Technologies) and 1% BSA (Sigma).

Fixed, permeablised, and blocked cells were then incubated with primary antibodies at a concentration of 1:500 unless otherwise stated. The following antibodies were used for experiments in this work: polyglutamylated tubulin (mouse monoclonal antibody, clone B3 T9822, Sigma), pan-polyglycylated antibody (mouse monoclonal antibody, AXO49, MABS276 Millipore), monoglycylated antibody (AXO 962 mouse monoclonal MABS277, EMD Millipore), kinesin-1 (rabbit monoclonal to KIF1B ab 167429, abcam), axonemal dynein, *β* tubulin (Rabbit polyclonal PA5-16863), tyrosinated tubulin (rabbit monoclonal antibody, clone YL1/2, MAB1864, EMD Millipore), acetylated tubulin (Lys40, 6-11B-1, mouse monoclonal antibody, Cell Signalling Technology). DNAL1 antibody (PA5-30643 Invitrogen).

After a 1 hour incubation in the relevant mix of primary antibodies. Cells were washed twice with PBS before incubation in secondary antibodies (Alexa-568-phalloidin, anti-rabbit Alexa-647, anti-mouse Alexa-588 (Life Technologies) for one hour at a dilution of 1:300 in PBS.

### Stem Cell Culture

Gibco human episomal induced pluripotent stem cell line was purchased from Thermo Scientific and cultured on Geltrex basement membrane in StemFlex medium (Thermo Scientific).

Routine passaging was performed using EDTA (Sigma), with single cell seeding performed for transfection and attempted clonal isolation through the use of TryplE (Thermo Scientific). Briefly, cells were washed twice with PBS and once with either EDTA (for clump passaging) or TrypLE (for single cell) before incubation in 1mL relevant detachment media for 3 minutes at 37°C.

For clump passaging, EDTA was removed and 1mL of StemFlex added. Cells were detached by triurtating media onto the bottom of the well and subsequently adding the required volume to fresh media (in a new, GelTrex coated plate).

For single cell seeding, TrypLE was diluted in 2mL StemFlex and the solution added to a 15mL falcon tube for centrifugation at 200g for 4 minutes. The supernatant was then discarded and the cell pellet resuspended in the required volume.

### iPSC MK differentiation

iPSC differentiation to mature, proplatelet forming megakaryocytes was performed using a protocol based on work published by Feng *et al*.(34). To summarise, cells were detached by clump passaging and seeded on dishes coated with Collagen Type IV (Advanced Biomatrix) at 5*µ*g/cm^2^. Cells were seeded overnight with RevitaCell (Life Technologies) to support survival on the new basement substrate. To begin the protocol cells were washed twice and incubated in phase I medium comprised of APELII medium (Stem Cell Technologies) supplemented with BMP-4 (Thermo Scientific), FGF-*β*, and VEGF (Stem Cell Technologies) at 50ng/mL each. Cells were incubated at 5% oxygen for the first four days of the protocol before being placed in a standard cell culture incubator for a further two days in freshly made phase I medium.

At day 6 of the protocol cells were incubated in phase II media comprised of APELII, TPO (25 ng/mL), SCF (25 ng/mL), Flt-3 (25ng/mL), Interleukin-3 (10ng/mL), Interleukin-6 (10 ng/mL) and Heparin (5 U/mL) (all supplied by Stem Cell Technologies).

Each day in phase II media suspension cells were spun down at 400g for 5 minutes and frozen in 10% FBS/DMSO. After 5 days of collection, all frozen cells were thawed for terminal differentiation. Terminal differentiation was performed by incubating cells in StemSpan II with heparin (5U/mL) and Stem Cell Technologies Megakaryocyte Expansion supplement on low attachment dishes (Corning).

### RNP Complexes

The IDT Alt-R®RNP system was used to target and knock-out *TUBB1*. crRNAs were ordered at 2nmol and resuspended in 20*µ*L TE buffer (IDT) for a final concentration of 100*µ*M. Atto-555 labelled tracrRNAs were ordered at 5nmol and resuspended in a volume of 50*µ*L for a final concentration of 100*µ*M.

To prepare small guide RNAs (sgRNA), equimolar ratios of both crRNA and tracrRNA were mixed with Nuclease Free Duplex Buffer (IDT). This mix was then incubated at 95°C for 5 minutes before allowing the reaction mix to cool at −1°C/second to 25°C. This mix was then spun down and complexed with either HiFi Cas9 V3 (1081058 - IDT) or Cas9 R10A nickase (1081063 - IDT) purified Alt-R®.

Cas9 protein was diluted to 6*µ*g per transfection and incubated with an equal volume of annealed sgRNA. This mix was left for 30 minutes at room temperature to form complete and stable RNP complexes.

### Stem Cell Transfection

iPSC transfection was performed using Lipofectamine Stem (Life Technologies) according to manufacturer instructions. Briefly, iPSC were seeded on 24 well dishes coated with Geltrex at 50,000 cells per well. After an overnight incubation in StemFlex with RevitaCell, cells were washed twice with PBS and once with OptiMem before incubation in OptiMem with RevitaCell.

RNP complexes were prepared as described in section and resuspended in 25*µ*L OptiMem per reaction. A Lipofectamine Stem master mix was prepared using 25*µ*L OptiMem and 2*µ*L Lipofectamine STEM per reaction (4*µ*L if a donor template is included). Equal volumes of both Lipofectamine and RNP mix were incubated to form lipofection complexes over a 10 minute incubation at room temperature. The final transfection mix was added to cells in OptiMem and left for 4 hours before the addition of StemFlex medium (and any relevant small molecules).

Measurement of iPSC transfection efficiency after treatment with Lipofectamine STEM and IDT RNP complexes was performed using manual cell counting in Evos acquired images (Phase contrast and fluorescence).

### Microscopy

Images were acquired using an Axio Observer 7 inverted epifluorescence microscope (Carl Zeiss) with Definite Focus 2 autofocus, 63x 1.4 NA oil immersion objective lens, Colibri 7 LED illumination source, Hammamatsu Flash 4 V2 sCMOS camera, Filter sets 38, 45HQ and 50 for Alexa488, Alexa568 and Alexa647 respectively and DIC optics. LED power and exposure time were chosen as appropriate for each set of samples but kept the same within each experiment. Using Zen 2.3 Pro software, five images were taken per replicate, either as individual planes (spread platelets) or representative Z-stacks (resting platelets). Images were prepared for presentation using Fiji (ImageJ). LUTs were adjusted for presentation purposes, and a rolling ball background subtstraction applied. Where Z-stacks are taken, images are presented as a maximum intensity projection.

### Image analysis

Image analysis was performed using a customed workflow. Briefly, the actin channel from resting and spread platelet images was used to train Ilastik pixel classifiers (approximately 6 images per condition) for segmentation based on this channel. This was incorporated into a KNIME workflow which would run images through the classifier to generated segmented binaries in which co-localisation and fluorescence intensity statistics were calculated (35–37). For the data presented in this manuscript, *M* 1_*diff*_ (a corrected Mander’s co-efficient to channel 1) was used to determine the co-localisation of PTMs to tubulin, and an *M* 2_*diff*_ value (corrected Mander’s co-efficient to channel 2) was used to calculate the co-localisation of motor proteins to PTMs of interest (38).

### Statistics

Statistical analysis was performed using GraphPad PRISM 7. Specifics of each test are detailed in the figure legends of the relevant figures. *P* values below 0.05 were considered significant.

### Quantitative Real Time PCR (qRT-PCR)

To determine whether the 13 mammalian TTLLs and 6 CCPs were expressed in iPSC-MKs at the different stages of differentiation (day 1, day 5 and day 5 +heparin) a qRT-PCR panel was developed using TaqMan technology and an ABI 7900 HT analyser (Applied Biosystems, Warrington, UK). RNA samples were isolated and reverse-transcribed and amplified with the relevant primers using SYBR-Green based technology (Power SYBR(r) Master Mix, Life Technologies). Total RNA was extracted from iPSC cells using the NucleoSpin RNA kit (Machery-Nagel) and cDNA was synthesized using the High-Capacity cDNA Reverse Transcription Kit (Life Technologies). qRT-PCR was performed on all the TTLL/CCP fragments generated from primers designed in supplementary figure 5 and the housekeeping control GAPDH (GAPDHFOR 5′ − *GAAGGTGAAGGTCGGAGT* − 3′ and GAPDHREV 5′*GAAGATGGTGATGGGATTTC* − 3′).

Each reaction was set up in triplicate including a non-template control. Expression was analysed using the CT method using D1 undifferentiated cells as a control. A full list of primer sequences has been uploaded as figureS6.

### *TUBB1* Homology Modelling

Homology models of TUBB1 WT and mutations were made using SWISS MODEL software (39–42), using the solved TUBB3 heterodimer as a template (PDB: 5IJ0 (43)). TUBB1 and TUBB3 share approximately 80% sequence identity, and the model created corresponds to residues 1-425 of TUBB1.

## Results

### Identification and initial characterisation of *TUBB1* variants in patients with inherited thrombocytopenia and platelet dysfunction

Using whole exome sequencing of patients recruited to the GAPP (Genotyping and Phenotyping of Platelets) study, two C-terminal *TUBB1* variants were identified in unrelated families presenting with macrothrombocytopenia (Figure1A). Affected individuals in Family A were found to be heterozygous for a C>T transition resulting in an arginine to tryptophan amino acid substitution (c.C1075T, p.R359W) in *TUBB1*. Individuals in this family also carry a *GFI1B* variant (p.Cys168Phe). Variants in both genes have been linked to thrombocytopenia, however only individuals A:1 and A:3, both of whom carry *TUBB1* variants, present with a macrothrombocytopenia (107 and 85 × 10^9^/L respectively). Individual A:2 carries the *GFI1B* variant but is wild type for *TUBB1* and presents with a normal platelet count (221 × 10^9^/L). This shows that the *TUBB1* variant is responsible for the macrothrombocytopenia, and not *GFI1B*. Interestingly, individuals A:1 and A:3 also present with significantly higher immature platelet fractions (IPFs) and mean platelet volumes (MPVs) when compared to their *TUBB1* WT relative (53.5% and 55.1% compared to 25.5%, MPV for A:1 and A:3 too large for measurement). This variation in count and platelet morphology suggest that the *TUBB1* variant is causative of the macrothrombocytopenia observed.

**Fig. 1.**
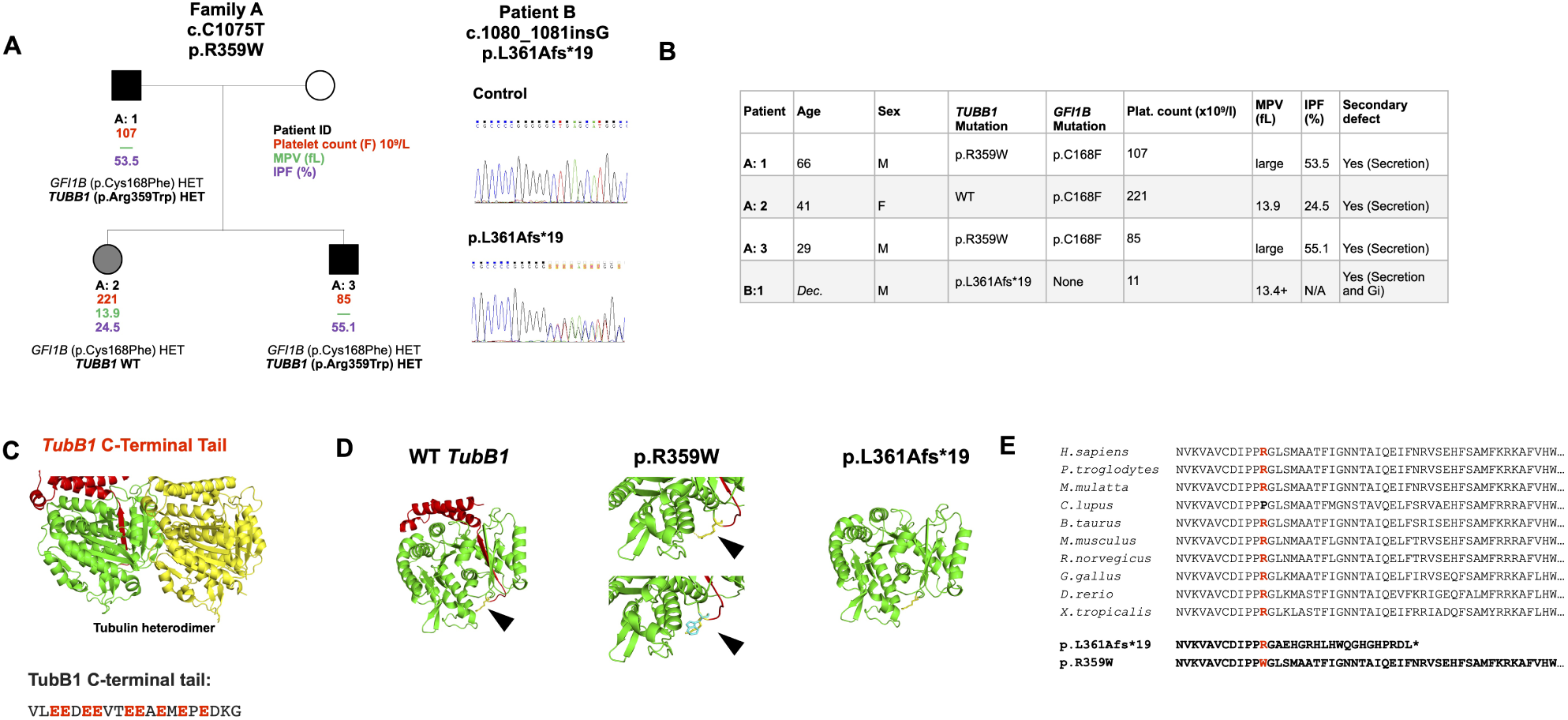
Candidate *TUBB1* mutations and their hypothesised effect on the C-terminus of *β*1 tubulin. (A,B) Two unrelated families were identified as carrying mutations in the *TUBB1* gene within 6 base pairs of one another. The first, family A, is comprised of 3 individuals, two of whom carry an Arginine to Tryptophan (p.R359W) coding mutation. Interestingly, all 3 individuals in family A harbour a *GFI1B* mutation. However, the individuals with the reported *TUBB1* mutation (A:1 and A:3) present with a macrothrombocytopenia and high IPF, while the patient without the R359W *TUBB1* mutation presented with a normal platelet count. The second family is comprised of a single individual, recently deceased, with a frameshift mutation 6 base pairs from the missense reported in family A. In this individual’s case, the insertion of a guanine nucleotide results in a frameshift with a premature stop codon 19 amino acids from the leucine to alanine change. (C) The C-terminal tail is downstream of both mutations in these families, and projects away from the dimer:dimer interface. The C-terminal sequence of *TUBB1* is rich in glutamate residues which can be targeted for polymodification. (D) Based on homology modelling of *TUBB1*, we predict that the missense mutation reported in family A is likely to affect the fold of the C-terminal tail, while the frameshift causes a truncation of this region. (Black arrows indicating the position of the R349 residue). (E) The arginine residue mutated in family A is highly conserved across species, as are sequences adjacent to the frameshift in patient B. (Note that the C-terminal tail of *TUBB1* is not presented in this model as there is no known or homologous structure available for this highly divergent sequence. The C-terminal portion of this model is used only as a means to visualise the effect of mutations on the C-terminal tail of *TubB1*.

Family/patient B was an elderly gentleman (now deceased) with a G insertion and subsequent frameshift truncation of the *β*1 tubulin protein 19 amino acids from the site of insertion (c.1080insG, p.L361Afs* 19). This patient had a severe macrothrombocytopenia with a platelet count of 11 × 10^9^/L and an MPV above 13.4 (Figure1B). At the time of study, IPF measurement was unavailable.

p.L361Afs* 19 is, absent in the latest version of the gnomad database, while *TUBB1*. p.R359W is a rare variant with a frequency of 6.78e x 10^−^3. Both *TUBB1* variants are positioned in the C-terminal region of the *β*1 tubulin as indicated in figures1C and D. This region is positioned away from the dimer interface with *α*-tubulin, and the C-terminal tail is an established site for PTM, particularly as it is rich in glutamate residues which are known targets for glutamylation and glycylation (1)(Figure 1C,D). Both affected *TUBB1* sequence variants are highly conserved in mammals (Figure1E). We predicted that the R359W missense substitution may alter the folding and positioning of the C-terminal tail as R359 forms direct polar contacts with the N-terminal helix holding the C-terminal region in place. The mutation to the hydrophobic tryptophan would destabilise the positioning of the C-terminal regions, potentially affecting PTM or interactions with critical microtubule accessory proteins (MAPs) (Figure1C,D). Similarly the G insertion and subsequent frameshift effectively deletes the C-terminus of the protein and would thus, if the protein folds correctly, have a similar predicted effect.

Patient platelet function was investigated using flow cytometry due to the reduced platelet count observed. Patient B demonstrated a significant reduction in surface P-selectin expression and fibrinogen binding in response to all agonists tested (Figure S1). Family A showed no change in the levels of surface receptor expression, but weak P-selectin and fibrinogen responses when activated with a low concentration ADP, CRP, and PAR-1, suggesting a mild secretion defect (FigureS1C,D).

Patients with C-terminal variants in this study and others previously reported by Fiore *et al*. phenocopy individuals with a complete loss of the *β*1 tubulin (18–20), suggesting that the C-terminal tail is likely critical to the function of *TUBB1* in the myriad complex roles of microtubules in both MKs and platelets. As this C-terminal tail is rich in glutamate residues which are often targeted for polymodification, we began to investigate the role of polyglutamylation and polyglycylation in human stem cell derived megakaryocytes and healthy donor platelets. Recently polyglutamylation of murine MKs and human platelets have been reported (30), however to date glycylation of these residues, and their mechanistic effects remain unknown.

### C-terminal mutations of *β*1 tubulin fold correctly and demonstrate a reduction in polymodification

First, we investigated the effects of the two patient mutations (p.R349W and p.L361Afs* 19) on the folding and potential polymodification of *β*1 tubulin. We designed, generated, and validated a *β*1 tubulin-mApple fusion over-expression construct (FigureS2). This plasmid was further mutated to harbour each of the patient mutations (p.R349W and p.L361Afs* 19), and an artificial C-terminal truncation which specifically deletes the glutamate rich C-terminal tail (FiguresS2,2A).

The wild type *β*1 tubulin construct was first transfected into Hek293T and co-stained for polyglutamylated and polyglycylated tubulin specifically. Interestingly, successfully transfected cells are exclusively positive for both polyglutamylated and polyglycylated tubulin indicating that *β*1 tubulin is indeed polymodified (Figure2B).

**Fig. 2.**
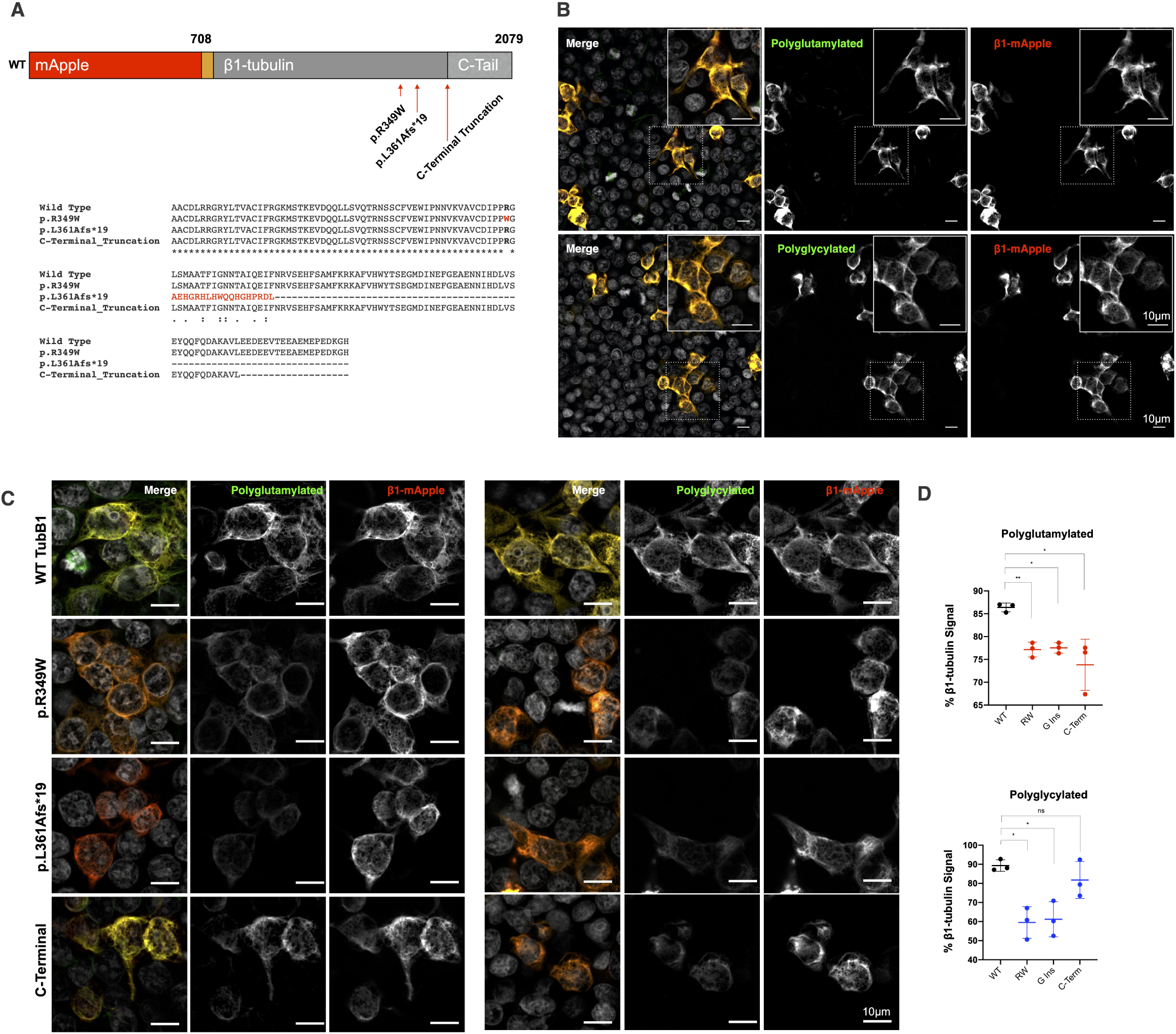
Cells expressing wild type and mutated *β*-tubulin demonstrate C-terminal polymodification. (A) Constructs carrying wild type, patient mutated (p.R349W and p.L361Afs* 19), and a C-terminal tail truncation of *β*1 tubulin fused to the fluorescent reporter mApple at the N-terminus were designed and cloned. (B) The wild type construct was first transfected into Hek293T cells, which were then fixed and immunostained for polyglutamylated and polyglycylated tubulin residues specifically. Transfected cells demonstrate both polygutamylation and polyglycylyation, unlike neighbouring transfected cells. (C,D) A comparison of the wild type and mutated constructs show a reduction in polyglutamylation and polyglycylation in each mutant when compared to the wild type. (*n* = 3 independent differentiations, S.D. plotted on graphs. 2-way ANOVA with multiple comparisons performed to establish significance.)

Expression of each of the mutated constructs show that both patient mutations and the C-terminal truncation fold correctly (Figure2C). Furthermore each of these mutations results in a consistent and significant reduction in polymodification (Figure 2D). This data indicates that both patient mutations result in a correctly folded *β*1 tubulin which phenocopies a trunctation of the C-terminus.

### iPSC-derived proplatelet forming MKs are both polyglycylated and polyglutamylated

The tubulin code posits that a highly lineage and species specific expression of modifying enzymes mediates cell specific PTMs. While our transfection data successfully indicates that *β*1 tubulin is indeed polymodified, and that each of our patient mutations result in the expression of a functional *β*1 tubulin with a C-terminal truncation, data from both human MKs and platelets is needed to dissect the role of polymodification in these cells.

Polyglutamylation has recently been reported in a modified CHO cell line and human platelets, however no evidence of this PTM has been reported in human MKs. A species dependent variation in the expression of modifying enzymes has been reported and discussed, therefore a human model is required to investigate the role of this polymodification in platelet production (25, 30, 44). To date, polyglycylation has not been reported in either platelets or megakaryocytes, however both modifications target the same glutamate residues, suggesting a potentially competitive mechanism by which these PTMs are applied to *β*1 tubulin.

To investigate polymodification in human megakaryocytes, we adapted a directed differentiation protocol previously reported by Feng *et al*. to generate large populations of mature, proplatelet forming cells (Supplementary FigureS3). Cells were differentiated and stained for CD42b as a marker for mature, and hence *TUBB1* expressing, MKs, and both polyglutamylated and polyglycylated tubulin. CD42b^+^ cells, including proplatelet forming cells, were also found to be positive for both polyglutamylated tubulin and polyglycylated tubulin (Figure3A), while neighbouring cells in the sample negative for CD42b did not demonstrate these polymodifications (Figure3B). Across multiple differentiations we consistently yielded a purity of approximately 50-60% CD42b^+^ cells (Figure3C), which on analysis are positive for both polyglutamylated and polyglycylated tubulin (Figure3D). Finally, 100% of proplatelet forming cells observed across replicates were positive for both polyglutamylated and polyglycylated tubulin (Figure3E). The presence of polyglutamylated and polyglycylated tubulin in these samples was further confirmed through western blotting of mature iPSC lysate from three independent differentiations (Figure3F).

**Fig. 3.**
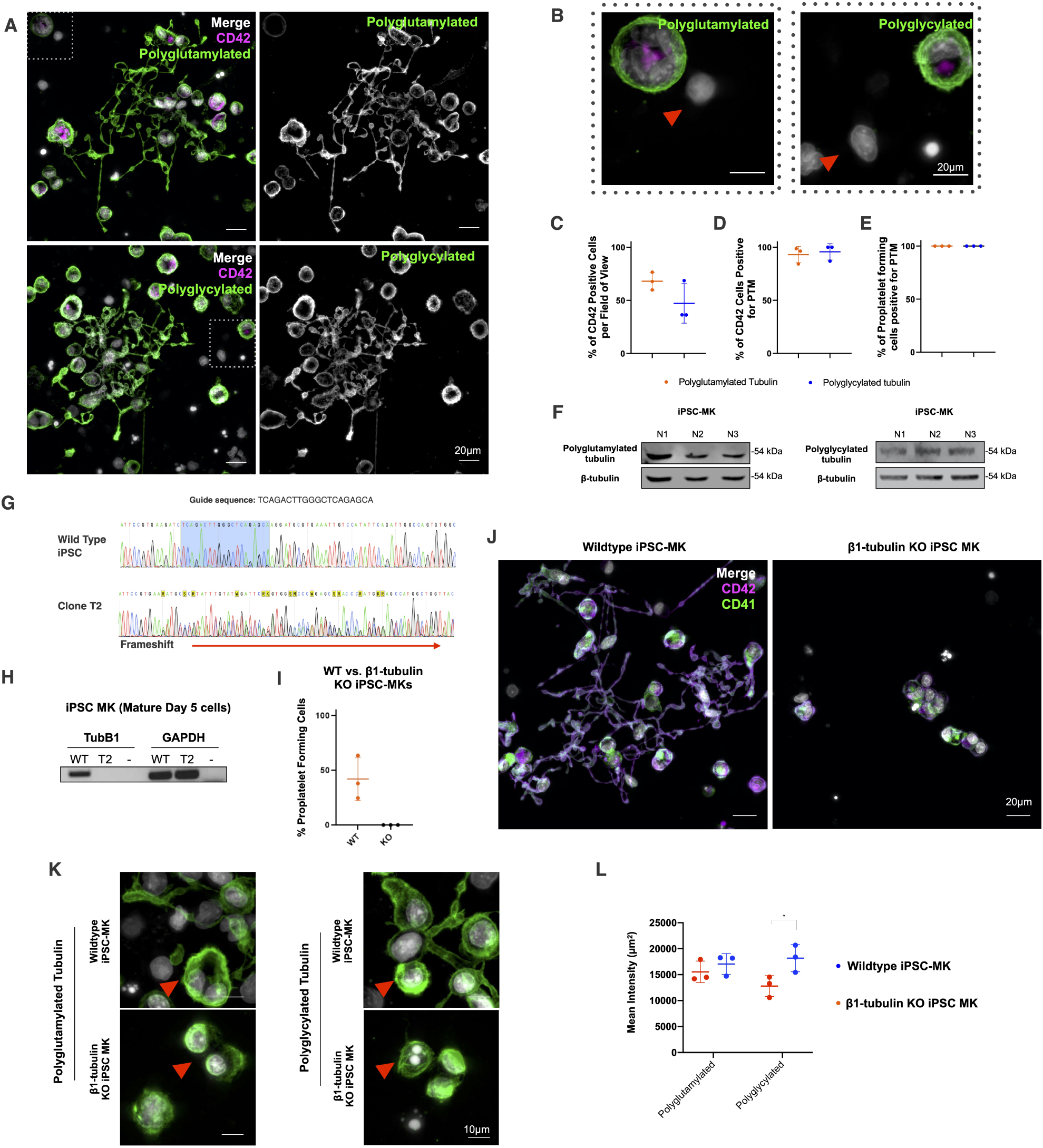
Mature, proplatelet forming iPSC-MKs are both polyglutamylated and polyglcylated. TubB1 knock-out iPSC-MKs do not form proplatelets and demonstrate disordered polymodified tubulin. (A) iPSC-MKs co-stained for CD42 and polyglycylated or polyglutamylated tubulin show that these cells are positive for both polymodifications. Both polyglutamylated and polyglycylated tubulin are evident in proplatelet extensions, including nascent platelet swellings on the proplatelet shaft. (B) Neighbouring CD42b^−^ cells are negative for both polymodifications (indicated by red arrows). (C,D) Approximately 50-60% of cells in multiple differentiations are CD42b^+^, and these cells are 100% double positive for polymodification and CD42b^+^. (E) All proplatelet extensions observed are positive for polymodification. (F) Polyglutamylation and polyglycylation are evident by western blotting in mature iPSC-MKs derived from 3 separate differentiations. (G) iPSC were transfected with a *TUBB1* targeting guide RNA, after which indel positive cells were isolated and sequenced to positively identify a bi-allelic insertion-deletion mutant (clone T2). (H) This clone was further analyzed and loss of *β*1 tubulin expression was confirmed by qRT-PCR. (I,J) A comparison of proplatelet production in wild-type vs. *TUBB1* KO cells revealed a complete loss of proplatelet formation in mutant CD41/42b^+^ cells. (K) *TUBB1* knock-out clones show a disordered arrangement of polymodified residues when compared to wild type cells. KO cells do not demonstrate the re-organisation of tubulin to the periphery of the cell evident in wild type iPSC-MKs (indicated by red arrows). (L) A measure of the polyglutamylation and polyglycylation in WT vs. KO clones reveals a significant increase in polyglycylation in mutant cells consistent with an aberrant accumulation of these residues. (*n* = 3 independent differentiations, S.D. plotted on graphs)

### CRISPR knock-out of *β*1 tubulin results in a complete loss of proplatelet formation

To date, the loss of *TUBB1* has not been studied in human MKs. To interrogate the loss of the protein, we generated an iPSC line with a CRISPR mediated biallelic loss of function mutation in the N-terminus of the coding region of *TUBB1* (Figure3G, Supplementary FigureS4). The mutation of the *TUBB1* start codon on both alleles results in a complete loss of protein expression (Figure3H), and a complete loss of platelet production *in vitro* (Figure3I,J). This is identical to what is observed in homozygous murine *TUBB1* knockouts (17). Unfortunately our attempts to generate C-terminal truncations through CRISPR in iPSC-MKs were unsuccessful as multiple guides targetting the 3’ end of the *TUBB1* gene failed to cleave the genomic sequence.

Interestingly while *TUBB1* knock-out clones stain positively for polyglutamylated and polyglycylated tubulin, the distribution of these residues is disturbed when compared to wild type platelet forming iPSC-MKs (Figure3K). While polyglycylated and polyglutamylated residues form a distinct peripheral band around wild type MKs as shown in figure3K, the knock-out cells demonstrate a diffuse tubulin staining.

This is the first data demonstrating the effects of the loss of *TUBB1* expression in human MKs, as well as the first report of both the polyglutamylation and polyglycylation of *β*1 tubulin.

### Platelet activation results in the polyglutamylation of the marginal band

MKs and platelets both achieve markedly different morphologies and functions despite the expression of *TUBB1* in both cell types. As such we hypotheized that the *β*1 tubulin polymodifications evident in MKs might be differently regulated between resting and activated platelets. We therefore compared immunofluorescence staining of polyglutamylated and polyglycylated tubulin between resting platelets and cells spread on fibrinogen and collagen.

We found that resting platelets demonstrate polyglutamylated tubulin which partially co-localises with the *β* tubulin marginal band (Figure4A). Unlike MKs which demonstrate extensive polyglycylation in proplatelet forming cells, platelets do not show polyglycylated tubulin (Figure4A).

**Fig. 4.**
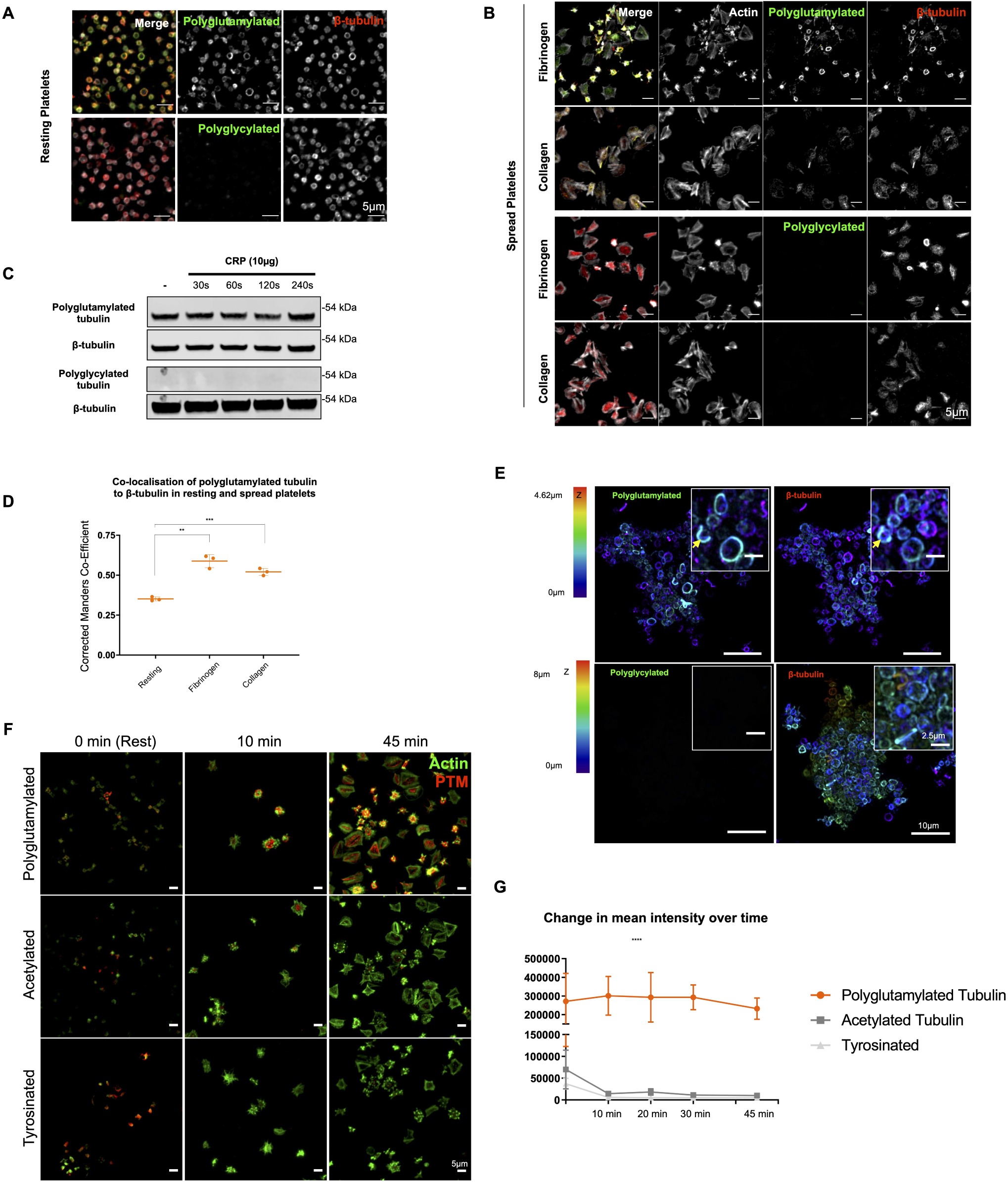
Platelet activation results in polyglutamylation of the marginal band. (A) Resting platelets show a partial polyglutamylation of the marginal band and a loss of the polyglycylation evident in iPSC-MKs. (B) Platelet spreading on fibrinogen and collagen shows an accumulation of polyglutamylation at the marginal band as platelets spread, and no evidence of polyglycylation. (C) Western blotting of resting and CRP activated platelets confirms the presence of polyglutamylated tubulin and a loss of polyglyclation. No increase in polyglutamylation is evident over time. (D) A measurement of co-localisation (corrected Manders coefficient) between polyglutamylated tubulin and *β*-tubulin in resting and spread platelets shows a significant increase in the co-localisation of these modified tubulin residues on platelet activation and spreading. (E) Platelets activated *in vitro* using Collagen Related Peptide (CRP) were fixed and co-stained for *β* tubulin and polyglutamylated residues, and then imaged in 3D using AiryScan confocal (stacks colourized in Z as indicated by the colour chart in this figure). In these micro-thrombi, polyglutamylation of the marginal band is evident, while polyglycylation is not observed. (F) A time course was performed to compare polyglutamylated tubulin with two other previously reported PTMs in platelets (acetylation and tyrosination). Polyglutamylation is significantly maintained over time, while acetylation and tyrosination decrease markedly as platelets spread. (G) The mean fluorescence intensity of polyglutamylated tubulin is markedly higher than either acetylated or tyrosinated tubulin. (*n* = 3, *S*.*D*., Two-Way ANOVA with multiple comparisons. *10 µm scale bar*.)

On fibrinogen and collagen spreading, polyglutamylation is evident, notably at the marginal band of spreading cells on fibrinogen (figure4B). A lack of polyglycylation is consistent in activated platelets.

Western blotting of resting platelets and cells activated through stimulation by CRP over time does not show an increase in the total amount of polyglutamylated tubulin in the sample (Figure4C). Interestingly, analysis of the co-localisation between *β* tubulin and polymodified residues shows a significant increase in co-localisation between polyglutamylated tubulin and */beta*1 tubulin (Figure4D). This suggests that while the total polyglutamylation of platelets remains unchanged throughout activation, the distribution of these residues is altered.

To investigate the distribution of both polyglutamylated and polyglycylated tubulin in the context of thrombi, platelets were activated *in vitro* using CRP and subsequently fixed and mounted on to Poly-L-Lysine coated coverslips. These cells were then stained for both *β* tubulin and polyglutamylated or polyglycylated tubulin respectively (Figure4E). Interestingly, polyglutamylated tubulin is evident throughout the aggregate on the marginal band of platelets, while polyglycylated tubulin is once again absent (Figure4F,G).

This data shows that while polyglutamylation and polyglycylation are evident in platelet producing iPSC-MKs, polyglycylation is completely lost in mature platelets. This is the first evidence of a system whereby these competitive polymodifications are dramatically altered between the ‘parent’ and terminal cell, suggesting markedly different roles for these residues in mediating the function and activity of */beta*1 tubulin.

While the total amount of polyglutamylated tubulin does not change on platelet activation, the localisation of these residues changes dramatically, suggesting that polyglutamylation of the marginal band specifically is key to the reorganisation of the microtubule cytoskeleton required for platelet function.

Acetylation and tyrosination have been previously reported in platelets, however their role in maintaining the marginal band and/or driving morphological change on platelet activation remains unclear(13). To determine whether the polyglutamylation of the marginal band we observe thus far coincides with these PTMs, we performed a time course of spreading on fibrinogen to determine whether there is an equivalent increase in either acetylation or tyrosination of the marginal band. Interestingly, we find a decrease in acetylation and tyrosination over time, while a notable polyglutamylation of the marginal band is evident from the earliest time point (10 minutes spreading on fibrinogen) (Figure4F,G).

### Platelet and MK polymodifications regulate motor protein localisation to drive both proplatelet formation and platelet shape change on activation

Thus far we have observed a markedly different distribution of polyglutamylated and polyglycylated residues in both human iPSC-derived MKs and human donor peripheral blood platelets. Polyglutamylation has been reported as a means by which motor protein processivity is regulated, and like in neuronal cells, MK proplatelet formation is known to be driven by a mechanism of dynein mediated proplatelet sliding (45). Similarly, the antagonistic movement of dynein and kinesin are known to maintain the marginal band in resting platelets(13).

We hypothesised that polyglutamylation in both MKs and platelets increases motor protein processivity and therefore alters the localisation of these proteins. Therefore the polyglycylation evident in MKs (but notably absent in platelets) is likely a mechanism of regulating motor protein motility to prevent excessive polyglutamylation and control the microtubule sliding required for platelet production. Interestingly, the polyglutamylation and polyglycylation of proplatelet extensions is analogous to the polymodifications observed in ciliated cells, in which axonema dynein also mediates the function of these unique structures.

To test this hypothesis, we first performed a time course of platelet spreading on fibrinogen and measured co-localisation between polyglutamylated tubulin and dynein (DNAL1 - axonemal light chain 1). We observe a sharp loss of co-localisation between dynein and polyglutamylated residues upon platelet spreading (Figure5A,B). Interestingly, axonemal dynein is also localised towards the leading edge of platelets when spread on fibrinogen (Figure5A). This data suggests that the increased polyglutamylation of the marginal band observed on platelet spreading drives an outward movement of axonemal dynein. To confirm the role of axonemal dynein specifically in this process, we also stained spreading platelets for cytoplasmic dynein and observe a central distribution, suggesting an alternative role for cytoplasmic dynein in platelets (FigureS5).

**Fig. 5.**
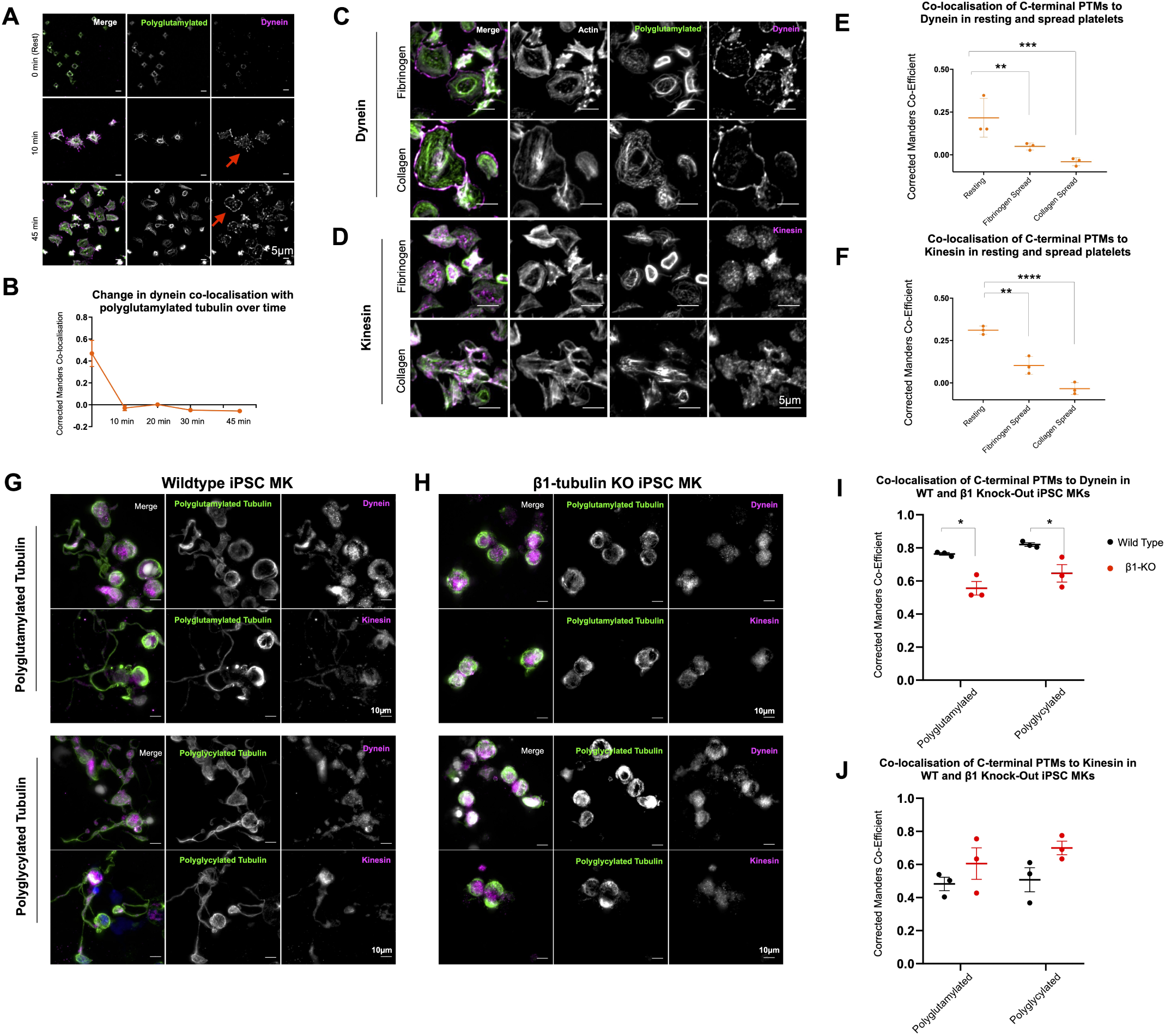
Polyglutamylation regulates spatial distribution of motor proteins in platelets and MKs. (A, B) Quantification of the co-localisation of dynein, as measured by the corrected Manders coefficient, with polyglutamylated tubulin over time shows a marked decrease in co-localisation between these polymodified residues and the motor over time. (C) In fibrinogen and collagen spread cells, dynein is observed on the periphery of the spread cells, (D) while kinesin-1 is evident as a diffuse punctate stain. (E) Dynein co-localisation with polyglutamylated residues decreases dramatically on platelet spreading in both fibrinogen and collagen (* * p = 0.0082 and p = * * * 0.0004 respectively). (F) a loss of co-localisation with polyglutamylated tubulin is evident in fibrinogen and collagen spread cells (* * p = 0.0017, * * * * p < 0.0001 respectively). (G) Immunofluorescence staining of iPSC-MKs for polymodified tubulin, dynein, and kinesin-1 show the distribution of both motors along the length of the proplatelet shaft in wild type cells. (H) In *β*1-tubulin knock-out iPSC-MKs, no proplatelet extensions are formed and a significant reduction in the co-localisation of dynein to polyglutamlated residues is observed (* p = 0.0166 and * p = 0.0293 respectively), (H,I) with no significant change in the co-localisation of kinesin-1 with polyglycylated tubulin. *n = 3, S*.*D*., *Two-Way ANOVA with multiple comparisons*.

We repeated a measure of co-localisation between polyglutamylated tubulin and dynein on both collagen and fibrinogen spread platelets at 45 minutes and observed a significant reduction in co-localisation, and a distinctive distribution of this motor protein at the periphery of fully spread cells on either substrate (5C, E). We repeated this experiment to investigate the effect of polyglutamlyation on the distribution of kinesin-1, a motor protein recently reported to be important in platelet secretion (46). A significant decrease in co-localisation is observed in both the fibrinogen and collagen spread cells (Figure5D, F).

This data suggests that polyglutamylated tubulin is involved in the localising motor proteins during platelet activation and spreading. The actions of kinesin and dynein are known to be critical to driving platelet shape change on activation, and this work is consistent with previous reports of polyglutamylated tubulin altering the processivity of motors in axons.

We then investigated the role of these polymodifications on the distribution of motors in iPSC-MKs. In proplatelet extensions axonemal dynein and kinesin-1 are both evident along the length of the proplatelet shaft (Figure5G). A mechanism of dynein mediated sliding has been reported as the driver of proplatelet elongation, however the original work cites cytoplasmic dynein as the mediator of this effect(45). Evidence of axonemal dynein in both the platelet and the proplatelet extension suggests that axonemal dynein likely plays a role in this process. *β*1 tubulin knock-out cells show no proplatelet formation, and a significant reduction in the co-localisation of dynein with polyglutamyulated residues when compared to wild type iPSC-MKs (Figure 5H,I). No significant change in the colocalisation of these residues with kinesin-1 is observed between wild type and knock-out iPSC-MKs.

### MK and platelet polymodification is regulated through the expression of both modifying and reversing enzymes

Thus far we have reported a mechanism by which polyglutamylation and polyglycylation occur in mature and proplatelet forming MKs, followed by a change in the distribution of these polymodifications (a loss of polyglycylation) in the resting platelet, and finally an increase in the polyglutamylation of the marginal band on platelet activation and spreading. We hypothesise that the expression of cell specific subsets of effecting (TTLL) and reversing (CCP) enzymes are required to achieve the observed regulation of these polymodifications. To date the expression of these enzymes in MKs and platelets has not been reported.

We designed a qRT-PCR panel to interrogate the expression of the 13 known mammalian TTLLs and 6 CCPs. We generated RNA from iPSC-MKs at different stages of the final terminal differentiation (Figures6A,S3A). Day 1 (d1) cells are representative of a pool of haematopoeitic stem cells (HSCs) and MK progenitors, while day 5 (d5) cells are comprised of 60% CD41/42b+ cells (Figures6A,S3E). Finally, day 5 cells treated with heparin to induce proplatelet formation (d5 + Hep) were used to interrogate whether there is any specific up-regulation of TTLLs and/or CCPs on proplatelet formation (Figures6A, S3D).

**Fig. 6.**
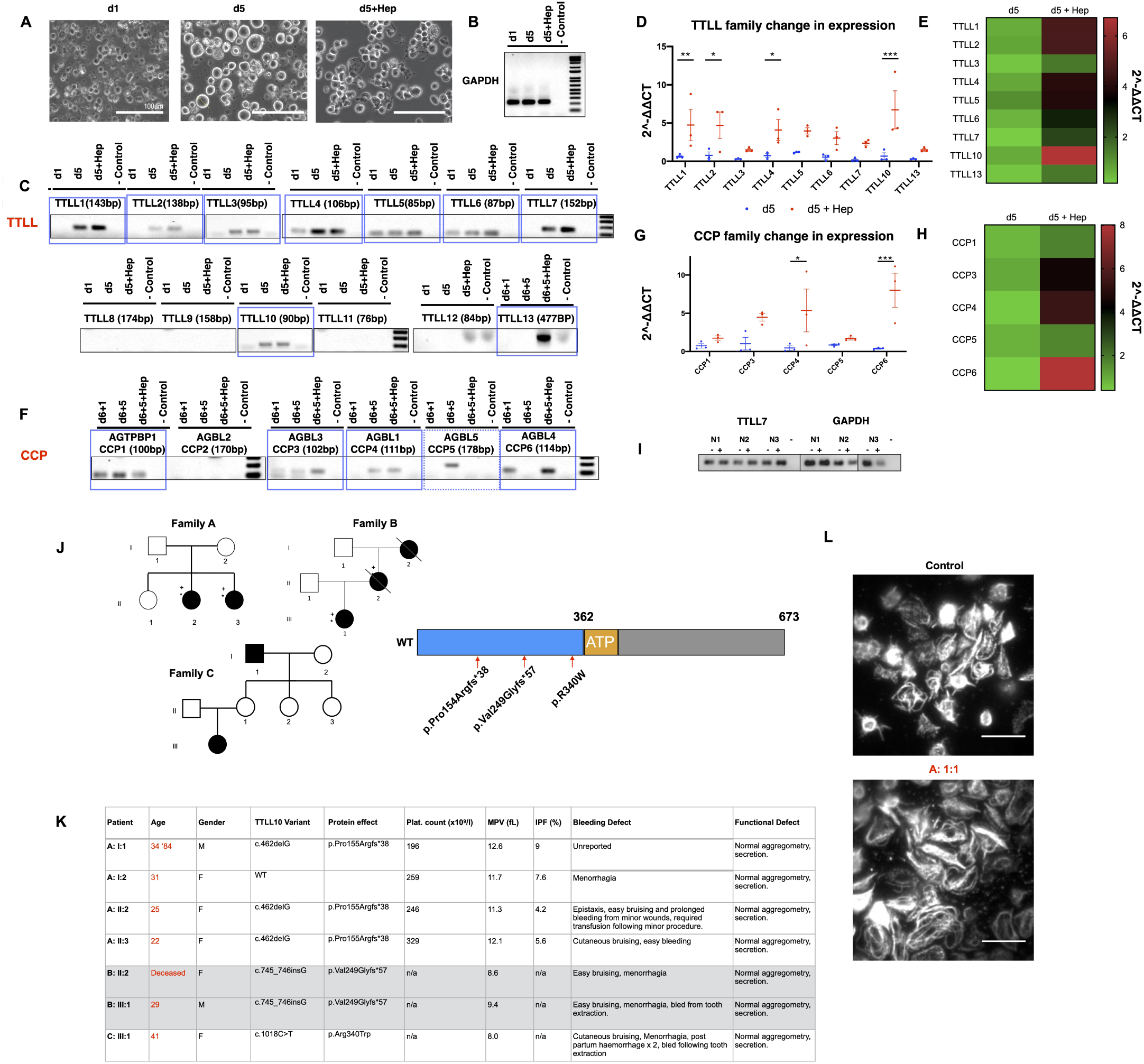
iPSC-MKs and platelets differentially express TTLL and CCP enzymes to regulate polymodifications during MK maturation and platelet production. The loss of TTLL10 results in a bleeding phenotype in unrelated human patients. (A) RNA was generated from iPSC-MKs at different stages of terminal differentiation. Day 1 (d1) cells are comprised of HSCs and progenitors, while day 5 (d5) cells are primarily mature CD41/CD42b double positive cells. Finally, day 5 cells treated with heparin to induce proplatelet formation (d5 + Hep) were used to determine whether an upregulation of these enzymes is evident on platelet production. (B) Samples were amplified with housekeeping GAPDH primers to ensure that any differences in TTLL or CCP expression observed were not due to differences in RNA abundance or quality. (C) A number of TTLL family enzymes were observed, including TTLLs1, 2, 3, 4, 5, 6, 7, 10, and 13. Of these, a number appeared to be up-regulated in mature and proplatelet forming cells. (D) These samples were taken forward and expression was quantified over multiple differentiations using the *dd*CT method using d1 cells as controls to determine if there was any upregulation in TTLL expression on platelet production. TTLL 1, 2, 4, and 10 expression was found to be significantly upregulated on treatment with heparin (* * p = 0.0081, * p = 0.0105, * p = 0.0260, * * * p = 0.0004 respectively). (E) A similar panel was performed on CCP enzymes which reverse polymodifications, with expression of CCP1, 3, 4, 5, and 6 was observed. (F) Statistically significant upregulation of CCPs 4, and 6 were observed on proplatelet production (* p = 0.0130, * * * p = 0.0009). (G) In resting (-) and CRP activated (+) platelets from 3 healthy donors TTLL7 was the only modifying enzyme found to be consistently expressed. (J) Three unrelated families were identified within the GAPP cohort, two with frameshift truncations and one with a missense (p.Pro15Argfs* 38, p.Val249Glyfs* 57, and p.Arg340Trp respectively). (K) All three families present with normal platelet counts and function, but demonstrate an elevated mean platelet volume (MPV) and consistent histories of bleeding including cutaneous bruising and menorrhagia. (L) Patient A:1:1 volunteered to return and demonstrated abnormally large platelets on spreading on fibrinogen (tubulin staining, 10 *µ*m scale bar). (*n* = 3, S.D., Two-Way ANOVA with multiple comparisons performed. Complete unedited gels found in figures S7,S8.)

GAPDH housekeeping controls for each of the 3 samples (d1, d5, d5+Hep) show equivalent amplification of the housekeeping control, while results for TTLL family proteins show a number of enzymes expressed at different levels across the maturation of these cells (Figure6B,C). Candidate TTLLs observed in the initial endpoint PCR were taken forward for quantification across replicates generated from multiple differentiations, which included TTLLs 1, 2, 3, 4, 5, 6, 7, 10, and 13 (Figure6C,D). We find a significantly increased expression of TTLL1, TTLL2, TTLL4, and TTLL10 on proplatelet formation in cells treated with heparin (* * p = 0.0081, * p = 0.0105, * p = 0.0260, * * * p = 0.0004 respectively) (Figure6D).

CCP family enzymes were also found to be expressed in maturing MKs, notably CCP1, 3, 4, 5, and 6 (Figure6E). Quantification across replicates revealed that CCP4 and 6 are up-regulated on proplatelet production (* p = 0.0130, * * * p = 0.0009) (Figure6F).

To determine whether platelets demonstrate a difference in TTLL and CCP expression we repeated the qRT-PCR panel performed on maturing iPSC-MKs on platelet samples from 3 healthy donors. Each donor’s platelets were either lysed in the resting state, or treated with CRP for 3 minutes, for RNA, to determine if there are changes in expression as a result of platelet activation. Interestingly, we found that none of the TTLLs and CCPs observed in iPSC MKs were consistently expressed across donors with the exception of TTLL7, a known polyglutamylase (Figure6I, complete gel in figureS8). No differences between resting and activated platelets were observed (Figure6I). This data shows a markedly different pattern of TTLL and CCP expression in both MKs and platelets, correlating with the observed differences in polymodification.

### *TTLL10* variants cause higher MPV and moderate to severe bleeding in 3 unrelated families

Our qRT-PCR screen reveals that a number of TTLL and CCPs are up-regulated during the process of platelet production, including the monoglycylase TTLL3 and the polyglycylase TTLL10. Three separate families were identified within the GAPP patient cohort with rare variants in the *TTLL10* gene (Figure6J). Two of the three variants result in frameshifts towards the N-terminus of the protein, preceding the ATP binding region (p.Pro15Argfs* 38 and p.Val249Glyfs* 57) (Figure6J). The final family has a missense p.Arg340Trp variant. p.Pro15Argfs* 38 is a novel variant, while p.Val249Glyfs* 57 and p.Arg340Trp are rare variants with a gnomad frequency of 2.15 × 10^−3^ and 3.34 × 10^−5^ respectively.

All three families report similar phenotypes, namely normal platelet counts, aggregation and secretion and an established history of moderate to severe bleeding, including cutaneous bruising and menorrhagia (Figure6K). Family A demonstrate a consistently consistently high MPV (normal ranges Mean Platelet Volume (fL) (7.83-10.5), while families B and C do not. Interestingly, one of the patients (A 1:1) was further studied, and on platelet spreading on fibrinogen coated coverslips we observed a marked increase in platelet area compared to controls (Figure6L).

## Discussion

Platelets and their progenitor cells, megakaryocytes, are a unique model for the study of the tubulin code. Like other specialist cells, they both express a lineage restricted isoform of *β* tubulin (*TUBB1*) which is linked to disease pathologies when lost (inherited macrothrombocytopenias)(5, 18–20). However, unlike other specialist cells which exemplify the tubulin code, MKs and platelets execute markedly different functions despite their shared *β*1 tubulin. MKs are the largest cell of the bone marrow, typified by a lobed, polyploid nucleus and the generation of long proplatelet extensions into the bone marrow sinusoids for the generation of platelets. Conversely platelets themselves are anucleate and the smallest circulating component of peripheral blood, classically involved in haemostasis and thrombosis, but with a myriad of other roles in wound healing, inflammation, and cancer progression.

A key question is therefore how *TUBB1*, restricted to both cell types, helps these cells achieve markedly different morphologies and functions. We hypothesised that a system of polymodification (polyglutamylation and polyglycylation) targeting the glutamate rich C-terminus of this *β* tubulin isoform, analogous to similar PTMs demonstrated in cilia and neuronal cells, is a likely mechanism by which the interactions of *β*1 tubulin with key motors are regulated in both MKs and platelets.

The study of this system is complicated by a lack of human cell lines which phenocopy primary MKs, namely in the generation of proplatelet extensions. Recent advances in the generation of MKs from iPSCs have allowed for the generation of large pools of mature CD41/42b^+^ cells, however few of these approaches have yielded large populations of MKs forming proplatelet networks equivalent to those produced by murine foetal liver cells (34, 47). In the study of the tubulin code and a system of PTMs, a species specific model is required due to controversies regarding the functionality of particular modifying enzymes. TTLL10 for example, has both been reported as redundant in humans by Rogowski *et al*., and as an ‘elongase’ requiring the expression of an initiating enzyme (TTLL3) to enact its function as a polyglycylase (25, 28,48). Therefore in this work we adapt an existing method of directed differentiation to produce large, proplatelet producing samples of iPSC-MKs for extensive immunfluorescence analysis of polymodifications and their association with motor proteins.

We report a system by which mature, CD42b^+^ iPSC-MKs demonstrate both polyglutamylation and polyglycylation. We do not observe any cells with proplatelet extensions lacking these polymodifications. Interestingly, we observe a markedly different distribution of these PTMs in the resting platelet, where polyglycylation is lost and polyglutamylation is partially co-localised to the marginal band. On platelet activation, we observe a marked change in the localisation of polyglutamylation specific to the marginal band. In iPSC-MKs with a CRISPR knock-out of *TUBB1*, we see a complete loss of proplatelet formation and lose the distinct re-organisation of polyglutamylated and polyglycylated tubulin around the periphery of MKs as seen in wild type cells.

MK proplatelet extensions are known to be driven by a system of dynein mediated microtubule sliding, while the marginal band in a resting platelet has been shown to be maintained by the antagonistic movement of dynein and kinesin (13, 14,45). Interestingly, polyglutamylation has been reported as a mechanism of altering motor protein processivity, with *in vitro* assays suggesting that polyglutamylation of *β* tubulin isoforms like *TUBB1* and *TUBB3* accelerates these motors (29). We show a significant effect of polyglutamylation on the spatial localisation of dynein and kinesin, supporting *in vitro* assays which suggest that polyglutamylation is an accelerator of motor proteins.

As both polyglutamylation and polyglycylation target the same substrate (a tubulin tail glutamate residue), it is likely that the competitive modification of these residues allows for the tight regulation of motor protein motility needed for proplatelet elongation. To investigate this, we report two unrelated patient families with C-terminal mutations (an R349W missense and a L361Afs* 19 causing macrothrombocytopenia, and show through the expression of these mutations in Hek293T cells that each mutation results in a functional *β*1 tubulin protein which phenocopies a C-terminal truncation.

The system of competitive polymodification evidenced in iPSC-MKs is analogous to the post-translational modification of ciliated cells, and so we reasoned that axonemal dynein, an isoform of the motor exclusive to axonemes, may play a role in both platelet formation and activation (6, 49,50). We find evidence of axonemal dynein on both proplatelet extensions and at the leading edge of spreading platelets. To our knowledge this is the first evidence of a functional role of axonemal dynein outside of classical ciliated structures. In our *TUBB1* knock-out MKs, we observe a decrease in the colocalisation of dynein to polyglutamyulated tubulin, suggesting that the loss of proplatelet formation observed in these cells is due to a dysregulation of the dynein-mediated microtubule sliding known to drive the elongation of the proplatelet shaft (45).

Our data suggests a tightly regulated, reversible system of polymodification which must be mediated by the cell specific expression of TTLLs and CCPs. Our expression profiles show a number of TTLLs and CCPs are expressed by MKs, while only the polyglutamylase TTLL7 is expressed by platelets consistent with our finding that in the platelet activation results in polyglutamylation of the marginal band, but no change in polyglycylation. In MKs we find an increase in expression of two TTLLs known to be involved in glycylation - the initiase TTLL3 and the elongase TTLL10. As previously mentioned, TTLL10 is the source of some controversy in the field, with reports suggesting the acquisition of a mutation in humans which renders the enzyme non-functional. Our findings support a role of TTLL10 as a polyglycylase on co-expression with TTLL3 as reported by Ikegami *et al*. in cell lines through co-transfection experiments.

In mice with a variant of the deglutamylase CCP1, an increase in polyglutamylation results in the degeneration of photoreceptors in the retina (51). A series of mutations in the human deglutamylase CCP5 have been reported in patients with visual impairments (52, 53). A loss of glycylation has similarly been reported to affect ciliary function and length in mice, and a loss of this PTM in photoreceptors results in ciliary shortening and subsequent retinal degeneration in mouse models (8, 54). A knock-out model of TTLL3 results in a loss of glycylation and the development of tumours in the colon (55).

In the absence of established TTLL and *TUBB1* specific inhibitors, the role of the PTM of *TUBB1* in human physiology is best understood by disease models, and much of our current understanding of the tubulin code is derived from correlating the loss of PTMs with human pathologies (5).

TTLLs are generally, ubiquitously expressed and extensively required for the development of neuronal, retinal, and ciliated cells. For the first time, we identify 3 unrelated families with *TTLL10* variants which result in an increase in platelet volume and an established history of bleeding which provides an invaluable insight to the potential role of polyglycylation in the context of platelet production and function. Our data shows that both TTLL3 and TTLL10 are expressed in platelet producing MKs. Our patient cohort do not lose TTLL3 function, and as such the action of TTLL3 as an initiase will occupy glutamate residues which would otherwise be polyglutamylated. In these patients we likely see a loss of polyglycylation, but no co-incident increase in polyglutamylation due to the normal function of the initiase (TTLL3). As polyglutamylation and monoglycylation are unaffected, platelet counts (and production) are normal, however, affected individuals appear to have an increased platelet volume and bleeding, suggesting a role for the extended glycine tail in regulating platelet size, with a downstream effect on the ability of platelets to prevent bleeding.

## Conclusions

The ‘tubulin code’ posits that a tightly regulated system of post-translational modification as well as the lineage restricted expression of tubulin isoforms is required to drive unique cellular behaviours. This has been shown through both human disease states and mouse models which demonstrate that the loss of cell specific isoforms or particular regulatory enzymes can result in a range of neuronal dysfunctions, ciliopathies, abnormal spermatogenesis (or sperm function), and platelet defects. The loss of *TUBB1* has been established as causative of macrothrombocytopenias, however the mechanisms by which *β*1 tubulin achieves the distinctive morphologies and functions of both MKs and platelets remains elusive.

Here we report a tightly regulated expression of glutamylating and glycylating enzymes through platelet production which drives the polyglutamylation and polyglycylation of MKs (Figure7). These modifications are reduced in the terminal platelet which only expresses the polyglutamylase TTLL7, and on activation the marginal band becomes heavily polyglutamylated to drive motor protein mediated shape change. We show a role for axonemal dynein in proplatelet extension and platelet spreading, and report novel variants of *TTLL10* which result in bleeding in patients through the loss of its’ role as a polyglycylase. Ultimately the role of this system of polymodification is to fine tune the motility of motor proteins in both the MK and the platelet, allowing both cell types to achieve their unique functions.

**Fig. 7.**
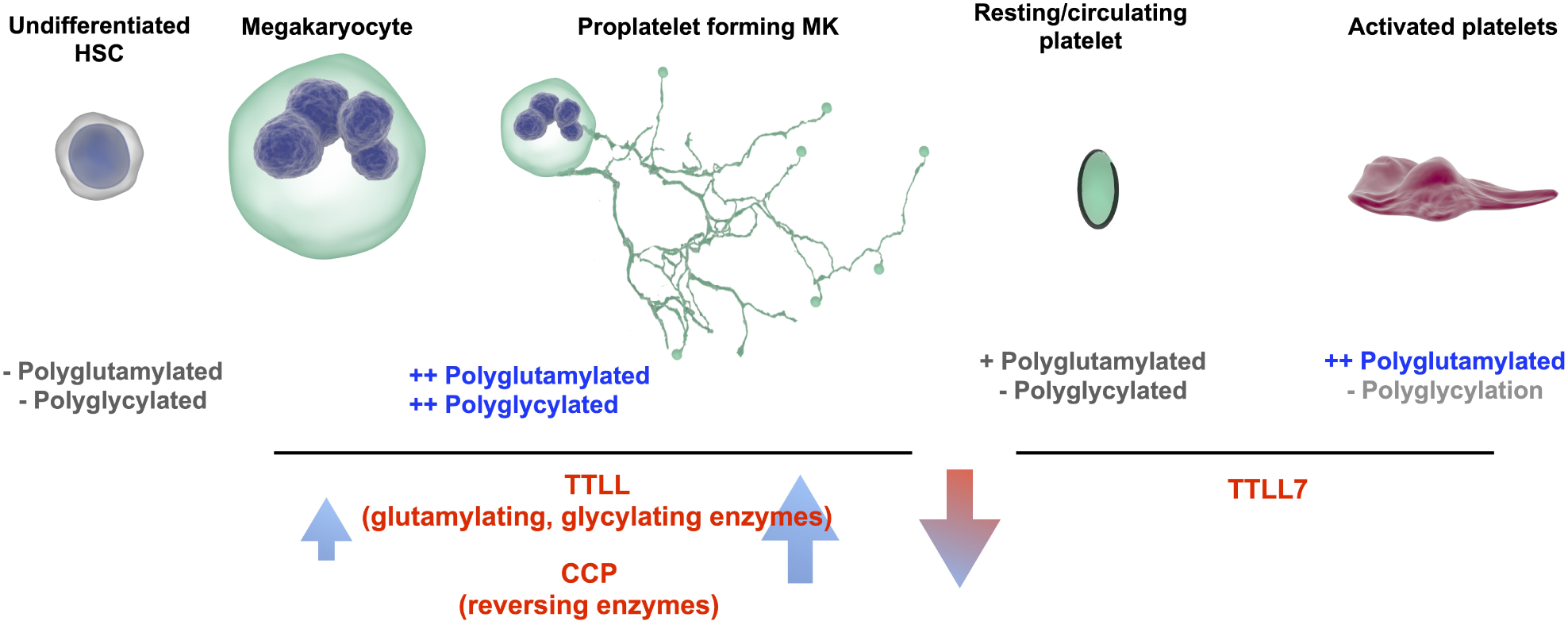
A system of competitive polymodification of *TUBB1* driven by the expression of TTLL and CCP enzymes is required for platelet production and function. We observe a system by which as iPSC-MKs mature and express *TUBB1*, they acquire both polyglutamylated and polyglycylated tubulin which co-incides with an increase in the expression of glutamylating and glycylating TTLLs and reversing CCPs. A resting platelet is partially polyglutamylated, and on activation the marginal band is polyglutamylated to drive shape change and spreading throguh the action of TTLL7. We show that this polymodification spatially affects the position of key motor proteins.

This work supports the paradigm of a ‘tubulin code’, and suggests that there is likely a complex regulatory system upstream of TTLLs and CCPs within an MK and a platelet which drives these PTMs.

## Author Contributions

AOK and NVM devised and performed experiments. AOK and NWM wrote the manuscript. AS performed homology modelling, cloning, transfection, and platelet spreading experiments. AM performed platelet spreading and analysis of patient sequencing data. PLR performed platelet preparations and reviewed the manuscript. JAP developed and applied image analysis workflows. JSR assisted with western blotting. JY contributed to platelet preparations. SGT, RS and NVM reviewed the manuscript.

## ACKNOWLEDGEMENTS

We thank the families for providing samples and our clinical and laboratory colleagues for their help. This work was supported by the British Heart Foundation (PG/13/36/30275; FS/13/70/30521; FS/15/18/31317; PG/16/103/32650). The authors would like to thank the TechHub and COMPARE Core facilities at the University of Birmingham. We thank Professor Steve Watson for his ongoing support and invaluable mentorship.

The authors do not have any conflict of interest.

