## Supplementary Material for "Post-translational polymodification of *β*1 tubulin regulates motor protein localisation in platelet production and function"

### Supplementary Figures and Tables

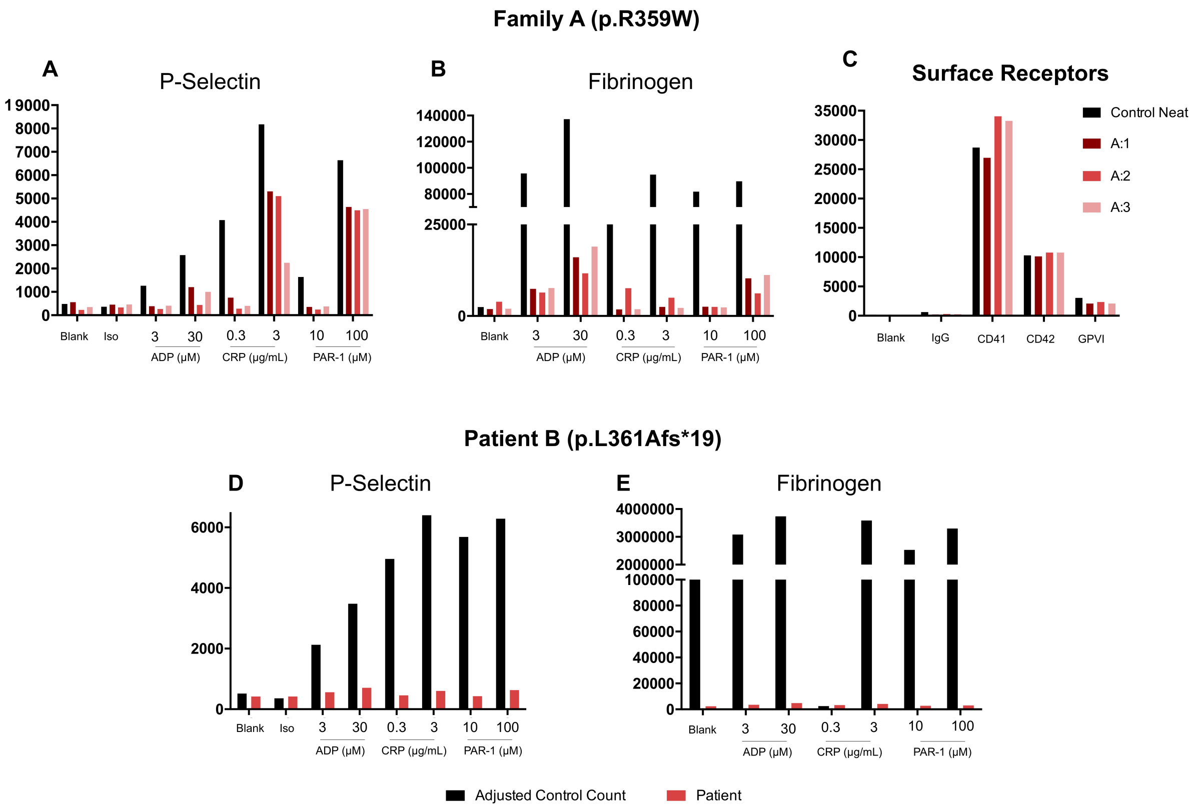


**Fig. S1. Patient flow cytometry data reveals secondary defects.** The GAPP project collects phenotypic data on patient recruitment, allowing for the assessment of secondary defects through FACS screening. (A) Individuals from family A show a reduction in P-selectin at both concentrations of CRP, low concentration CRP, and low concentration PAR-1. (B) Patients similarly show a reduction in fibrinogen uptake compared to controls, but show no difference in (C) surface marker expression. (D) Patient B shows a marked reduction in P-selectin surface expression and (E) fibrinogen uptake compared to controls.

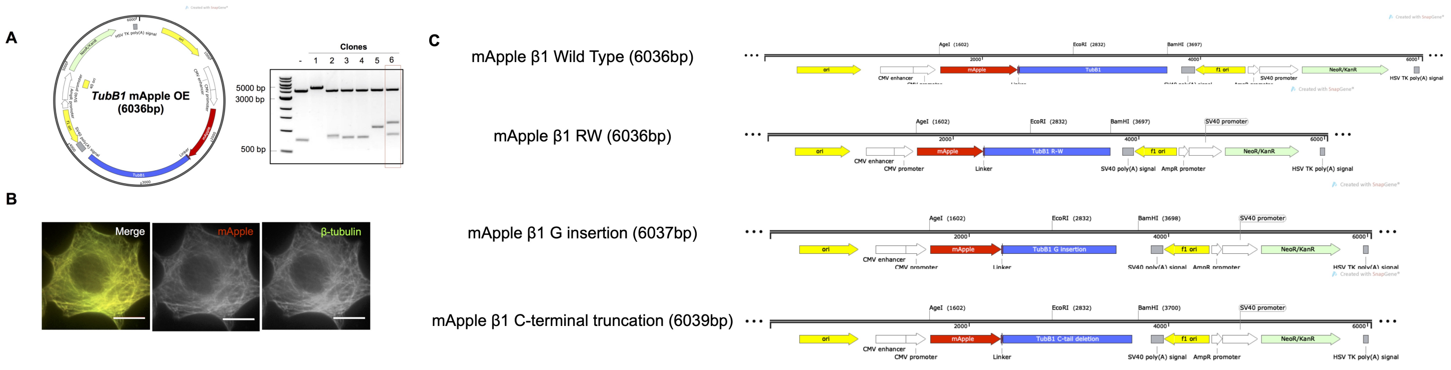


D

| | Primers | Annealing temperatures (°C) |
|---|---|---|
| **RW** | F: 5’ CATCCCGCCCtGGGGGCTGAG 3’<br>R: 5’ TCGCAGACAGCCACCTTGACG 3’ | 72 |
| **G insertion** | F: 5’ GCTGAGCATGGCCGCCACCTTC 3’<br>R: 5’ CCCCCGGGGCGGGATGTC 3’ | 72 |
| **C tail truncation** | F: 5’ TAAGAGGAAGATGAAGAGGTC 3’<br>R: 5’ TAGAACTGCTTTGGCATC 3’ | 59 |

**Fig. S2. Generation of wild type and mutated mApple-$\beta$1-tubulin plasmids.** (A) An N-terminal mApple-$\beta$1 tubulin over expression vector was designed and cloned through the gibson assembly of the $\beta$1 tubulin sequence into a C-terminal mApple construct (mApple-C1 was a gift from Michael Davidson (Addgene plasmid # 54631 ; http://n2t.net/addgene:54631 ; RRID:Addgene_54631). Of the 6 selected clones presented, clone 6 demonstrated cleavage bands of the predicted molecular weight, and was subsequently cloned. (B) The correctly assembled sequence was then transfected to and co-stained with a $\beta$-tubulin antibody to confirm the correct expression and fold of this tubulin construct. (C) Mutants of the wild type construct were generated through a Q5 site directed mutagenesis kit to generate constructs harbouring patient RW and G insertion mutants, as well as an artificially designed C-terminus truncation of the protein. (D) Primers used for the site directed mutagenesis are listed with their respective annealing temperatures.

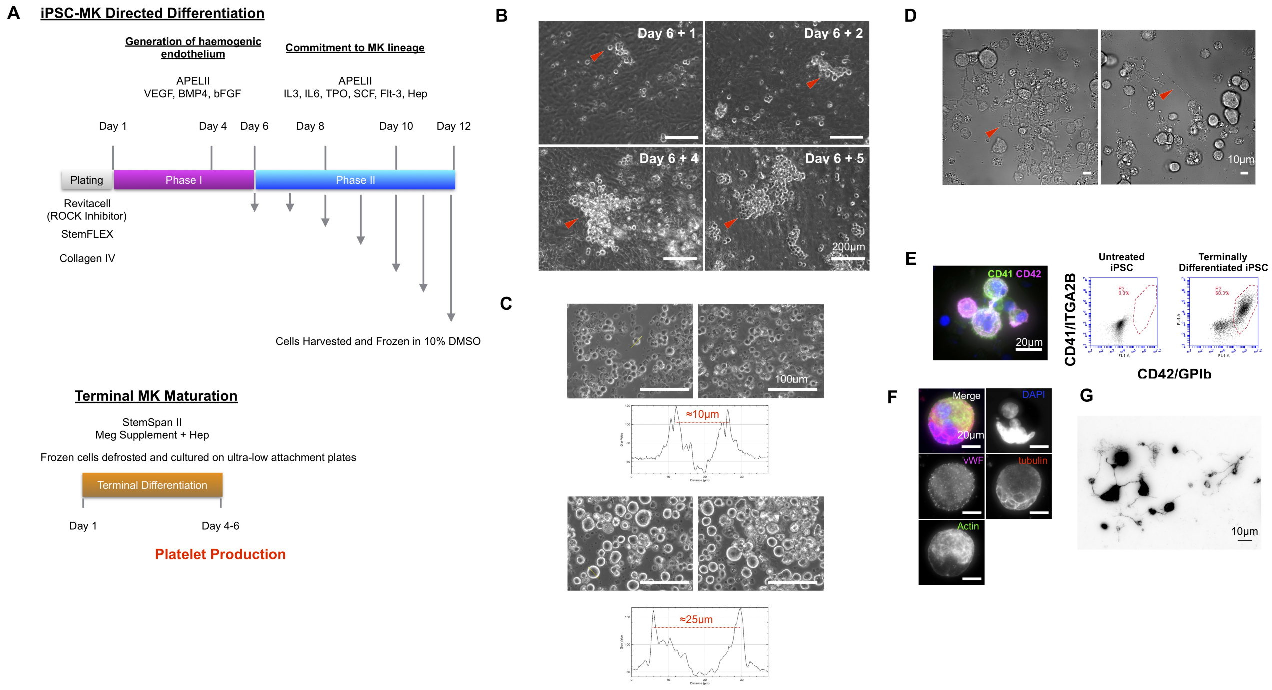


**Fig. S3. Directed differentiation of iPSC to proplatelet forming MKs.** (A) A 3 stage protocol was adapted from a method previously published by Feng *et al.*. Briefly, iPSC were clump passaged on to collagen IV coated plates and incubated in RevitaCell overnight before beginning Phase I of the differentiation. Phase I involves a 4 day incubation at 5% $O_2$ in APEL2 media supplemented with 50ng of BMP4, VEGF, and FGF2, after which fresh media was added and cells were incubated for 2 more days at normoxic conditions. Phase II of the protocol involved incubation in APEL2 supplemented with IL3, IL6, Flt-3, hSCF, TPO, heparin, during which time cells were harvested and frozen every 48 hours and fresh media added. Finally, harvested cells were thawed and incubated in StemSpan II medium with MK supplement for 5 days before samples were prepared for downstream assays (immunofluorescence, RT-PCR etc.). (B) During Phase II of the differentiation, progressively larger numbers of blast like cells are observed emerging from a layer of haemogenic endothelium. (C) During Phase III of the differentiation, cells grow from progenitors and blast like cells approximately 10 $\mu$m in size to large, mature MKs ranging in 25-40 $\mu$m in size. (D) At day 5, on treatment with Y-27632 and heparin, cells form elaborate proplatelet networks. (E) 60$\%$ of terminally (Phase III) differentiated cells are CD41 and CD42 double positive and on staining demonstrate (F,G) a mix of ploidies and proplatelet networks consistent with mature platelet producing MKs.

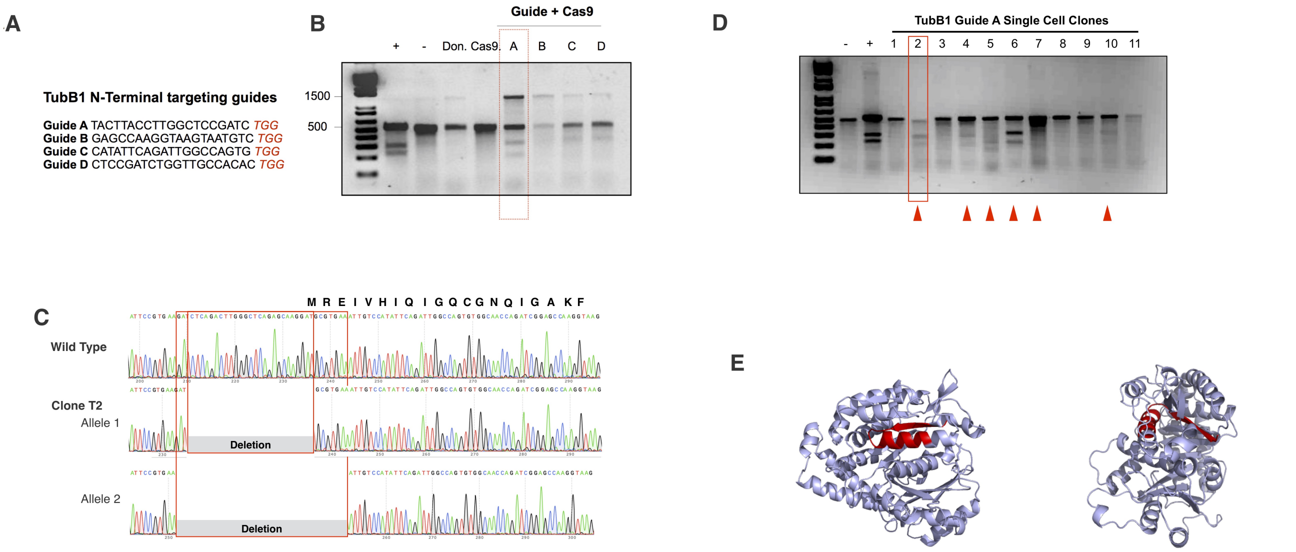


**Fig. S4. CRISPR bi-allelic knock-out of $\beta$1 tubulin.** (A) Guides targeting exon 1 of the *TUBB1* gene were designed and (B) tested for efficiency using a T7EI cleavage assay. The population evidencing the most cleavage (and hence most efficient guide (guide A)) was taken forward to generate *TUBB1* knock-out clones, through single cell clonal isolation. (C) Cells positive for cleavage on single cell expansion (identified by the red arrows) were taken forward for sequencing. (D) Clone T2 revealed a bi allelic loss of the start codon, (E) resuling in a deletion of a significant portion of the N-terminus and as evidenced in the main text, a loss of *TUBB1* expression.

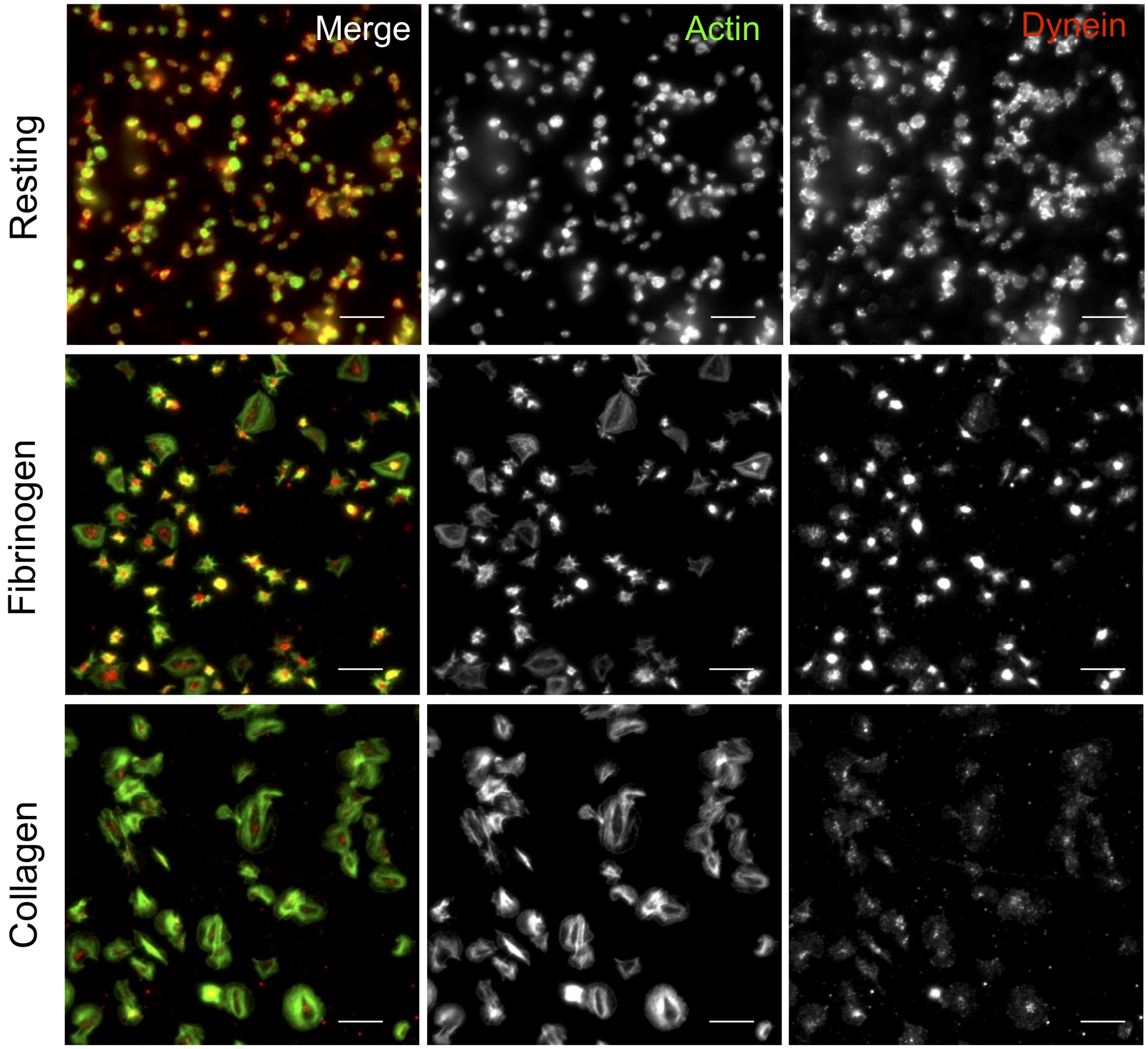


**Fig. S5. Staining of Cytoplasmic Dynein in resting and Spread Platelets.** Resting and spread human donor platelets were stained for cytoplasmic dynein to compare the distribution of this isoform of the motor protein to axonemal dynein. While axonemal dynein is primarily found on the edge of spreading cells, cytoplasmic dynein is found at the centre of spread cells, suggesting that the axonemal variant is involved in platelet spreading and activation.

| Name | Sequence : (5' to 3') | Fragment Size | Name | Sequence : (5' to 3') | Fragment Size |
|---|---|---|---|---|---|
| FH1_TTLL1 | AGTCAACCATTTTCCAAACC | 143 bp | FH1_TTLL11 | ATTTGTTTATCCGGTTCCTG | 76 bp |
| RH1_TTLL1 | AGTCCAGATAGAGGTATTTTCC | | RH1_TTLL11 | CTCCTTATGAAGGTACGAAAG | |
| FH1_TTLL2 | GCCTTTACCCTTAACATTCC | 138 bp | FH1_TTLL12 | CATTCTGGAGGAAAACAAGG | 84 bp |
| RH1_TTLL2 | TTTCTTCTTCTCCAGTGTTG | | RH1_TTLL12 | GTGTAGACCTTGAAGATGTG | |
| FH1_TTLL3 | AAGCCTTCATAGAGGACTTC | 95 bp | FH1_TTLL13 | ACCTGACCAACTATGCTATC | 477 bp |
| RH1_TTLL3 | TACTGCCTGAATAGGGTATG | | RH1_TTLL13 | TGGTTTTGATGATGATGTCC | |
| FH1_TTLL4 | GAAGCTAAACCATTTCCCAG | 106 bp | FH1_AGTPBP1 (CCP1) | AAAAACAAATGCCAGGAGAG | 100 bp |
| RH1_TTLL4 | GAAACTGAACTCCTTCTTGC | | FH1_AGTPBP1 (CCP1) | CATGTTTCTATGCCGGTTATC | |
| FH1_TTLL5 | AATTCATATTCGAAGGACCG | 85 bp | FH1_AGBL2 (CCP2) | GGCCTATCAGTTTATCTTCAG | 170 bp |
| RH1_TTLL5 | GATTGTTGATCAGGTAGACG | | RH1_AGBL2 (CCP2) | ATCTGTAATCCCAGCTACTC | |
| FH1_TTLL6 | AAGCCCTTTATCATTGATGG | 87 bp | FH1_AGBL3 (CCP3) | GAAGAGCAAAGAAGGAACAG | 102 bp |
| RH1_TTLL6 | GTACACAAAAATCCTGAGAGG | | RH1_AGBL3 (CCP3) | TTGTTACCCAGAGTAGATCC | |
| FH1_TTLL7 | CAGAATTGGTGGTAAAGACC | 152 bp | FH1_AGBL1 (CCP4) | AGATGATGACTTGGAAACAG | 111 bp |
| RH1_TTLL7 | CCATGGCTTTAGTTTTCTATCC | | RH1_AGBL1 (CCP4) | CTATAGGAGAGCTCAAGACAC | |
| FH1_TTLL8 | AACAAGGAATTTCCCAAGAC | 174 bp | FH1_AGBL5 (CCP5) | CTATATCCAAAGCTCATCTCC | 178 bp |
| RH1_TTLL8 | AGTGGAACTTCTTCTCTACC | | RH1_AGBL5 (CCP5) | AGTTGCATTCAAGTGTGTAG | |
| FH1_TTLL9 | ATCATGAAGCCTGTAGCC | 158 bp | FH1_AGBL4 (CCP6) | AAATGATGATGCCATTGGAG | 114 bp |
| RH1_TTLL9 | GGATTTTCAATGTAACGCTG | | RH1_AGBL4 (CCP6) | TTACCACTTTCAAAGCAAGC | |
| FH1_TTLL10 | GAAGAGTTTTTCCCAGAGAC | 90 bp | | | |
| RH1_TTLL10 | GATCCATATCTGGGTTTCATC | | | | |

**Fig. S6. Primers and predicted fragment lengths for qRT-PCR screen of TTLL and CCP expression.** Exon overlapping primers were disgned for a qRT-PCR screen of TTLL and CCP expression. Forward and reverse primer sequences are listed, along with predicted fragment length.

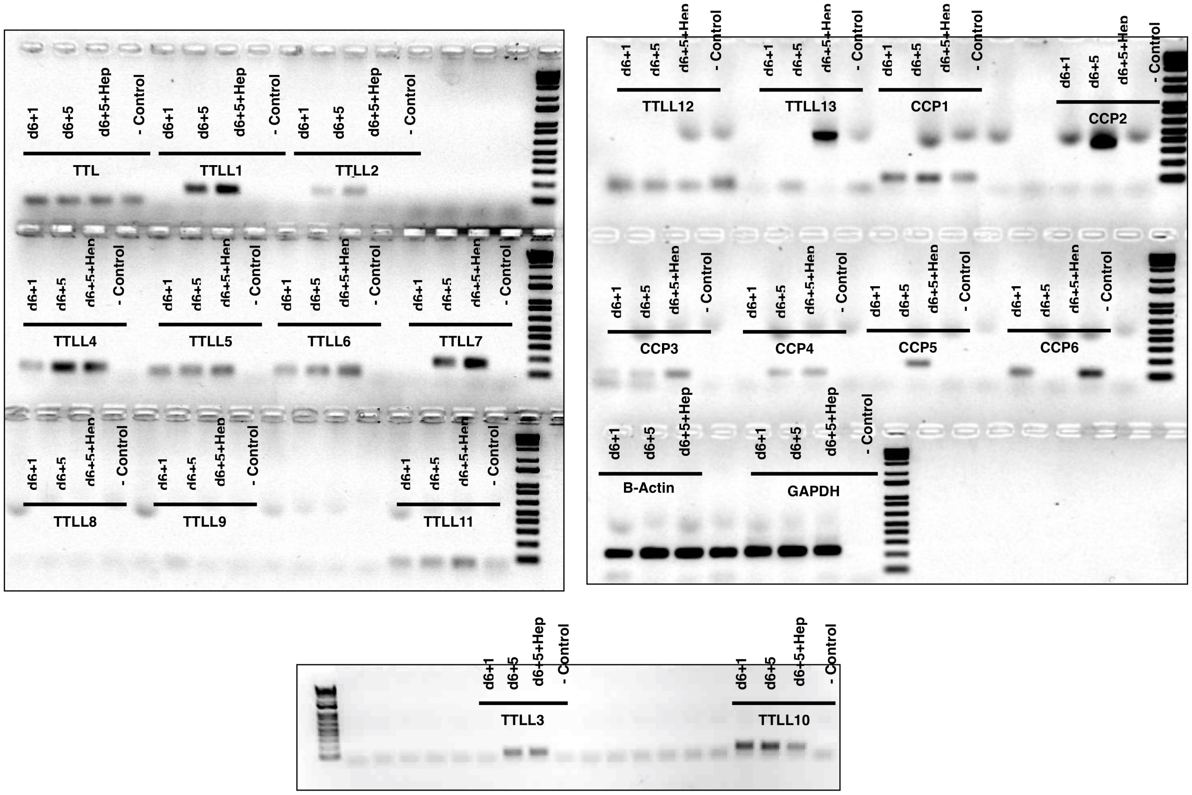


**Fig. S7. Whole gel for TTLL and CCP RT-PCR screen in iPSC-MKs.** Complete gels used in figure 6.

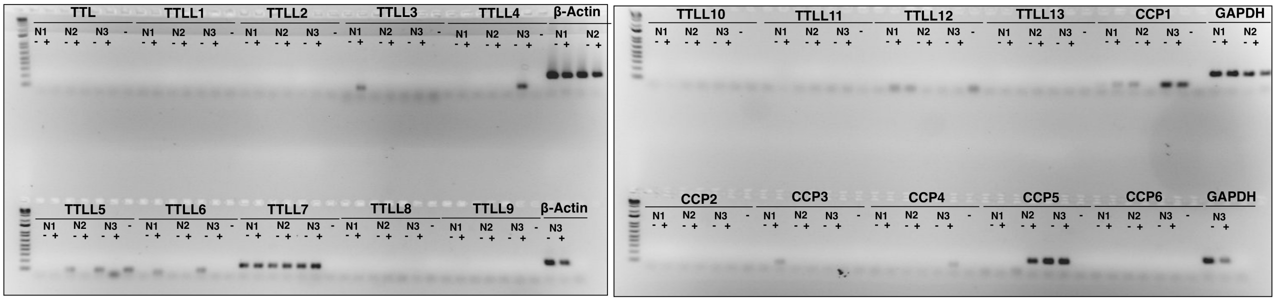


**Fig. S8. Whole gel for TTLL10 and CCP RT-PCR screen in human peripheral blood platelets.** Complete platelet gel used in figure 6.
